# The Hsp70/Hsp90 Co-Chaperone Hop/Stip1 Shifts the Proteostatic Balance from Folding Towards Degradation

**DOI:** 10.1101/562637

**Authors:** Kaushik Bhattacharya, Lorenz Weidenauer, Tania Morán Luengo, Pablo C. Echeverría, Lilia Bernasconi, Diana Wider, Matthieu Villemin, Christoph Bauer, Stefan G. D. Rüdiger, Manfredo Quadroni, Didier Picard

## Abstract

Hop/Stip1/Sti1 is thought to be essential as a co-chaperone to facilitate substrate transfer between the Hsp70 and Hsp90 molecular chaperones. Despite this proposed key function for protein folding and maturation, it is not essential in a number of eukaryotes and bacteria lack an ortholog. We set out to identify and to characterize its eukaryote-specific function. Human cell lines and the budding yeast with deletions of the Hop/Sti1 gene display reduced proteasome activity due to inefficient capping of the core particle with regulatory particles. Unexpectedly, knock-out cells are more proficient at preventing protein aggregation and at promoting protein refolding. Without the restraint by Hop, a more efficient folding activity of the prokaryote-like Hsp70/Hsp90 complex, which can also be demonstrated *in vitro*, compensates for the proteasomal defect and ensures an alternate proteostatic equilibrium. Thus, cells may act on Hop to shift the proteostatic balance between folding and degradation.

## INTRODUCTION

Homeostasis of the proteome, often referred to as proteostasis, is essential both at the cellular and organismic levels for health and longevity (Balch et al., 2008; Hipp et al., 2014; Labbadia and Morimoto, 2015). Proteotoxic stresses lead to protein misfolding and aggregation, which trigger and are counterbalanced by protein quality control mechanisms. The major cellular quality control mechanisms, all assisted by molecular chaperones, are protein refolding and the degradation of misfolded and aggregated proteins by the proteasome and by autophagy (Wong and Cuervo, 2010; Chen et al., 2011; Hipp et al., 2014; Balchin et al., 2016). Defects in any of these mechanisms can cause severe proteotoxicity, which in turn can lead to diseases such as cystic fibrosis, lysosomal storage diseases, cancer, and, most notably, neurodegenerative disorders such as Huntington’s, Parkinson’s, and Alzheimer’s diseases (Balch et al., 2008; Chen et al., 2011; Hipp et al., 2014; Schmidt and Finley, 2014; Labbadia and Morimoto, 2015).

In eukaryotes, the Hsp70 and Hsp90 molecular chaperone machines are major contributors to proteostasis by providing a platform for folding of both nascent polypeptides and misfolded, structurally labile and mutated proteins, collectively called “clients” (Mayer and Bukau, 2005; Echeverria and Picard, 2010; Kampinga and Craig, 2010; Picard, 2012; Taipale et al., 2012; Schopf et al., 2017; Radli and Rudiger, 2018; Moran Luengo et al., 2019). For folding and assembly of clients, both Hsp70 and Hsp90 undergo large conformational changes and collaborate with co-chaperones (Li et al., 2012; Mayer and Le Breton, 2015; Schopf et al., 2017). One of these co-chaperones is the Hsp70-Hsp90 organizing protein (Hop), encoded by the gene *STIP1* in mammals. It is an adaptor molecule between the Hsp70 and Hsp90 molecular chaperone machines, which facilitates the folding or stabilization of clients by promoting their transfer from the Hsp70 to the Hsp90 molecular chaperone machines after the initial recognition and binding of clients by Hsp70 in collaboration with its J-domain containing co-chaperone Hsp40 (Scheufler et al., 2000; Kirschke et al., 2014; Mayer and Le Breton, 2015). Hop forms a ternary complex with Hsp70 and Hsp90 using its tetratricopeptide repeat (TPR) domains. Two of its three TPRs, TPR1 and TPR2A, specifically bind the extreme C-terminal peptide sequences EEVD and MEEVD of Hsp70 and Hsp90, respectively (Scheufler et al., 2000; Schmid et al., 2012; Bhattacharya et al., 2018).

Proteins, whose folding or refolding fails, either because they cannot fold by themselves or with the assistance of molecular chaperones, are degraded by the proteasome, a highly conserved and regulated eukaryotic protease complex. About 80% of total cellular protein turnover is through this complex (Collins and Goldberg, 2017); moreover, the proteasome works together with an Hsp40-Hsp70-Hsp110 protein disaggregase complex to eliminate intracellular aggregates (Shorter, 2011; Hjerpe et al., 2016). The proteasome is a 1.6 to 2.5 MDa complex consisting of a 20S proteolytic core particle (CP) and a 19S regulatory particle (RP); the CP can be capped by one or two RPs resulting in 26S or 30S particles, respectively (Murata et al., 2009; Gallastegui and Groll, 2010). The RP is divided into a lid and a base and has unique regulatory functions; it recognizes ubiquitinated substrates produced by the E1-E2-E3 ubiquitination system, promotes their deubiquitination and unfolding and the subsequent gate-opening of the CP, and finally the loading of the processed substrates into the proteolytic chamber (Collins and Goldberg, 2017). Dedicated chaperones for the assembly of CP and the RP base are well known, whereas RP lid assembly is still not well understood (Murata et al., 2009). Interestingly, Hsp90 has been proposed to be an assembly chaperone for the RP lid complex based on genetic interactions in the budding yeast (Imai et al., 2003) and the reconstitution of the RP lid complex in *E. coli* co-expressing yeast Hsp90 (Lander et al., 2012).

Prokaryotes do have Hsp70 and Hsp90 orthologs but lack a Hop-like protein. Bacterial Hsp90 and Hsp70 physically and functionally interact directly without a Hop-like protein (Genest et al., 2015; Kravats et al., 2017). In eukaryotes, Hop is not absolutely indispensable as mutant budding yeast, worms (*Caenorhabditis elegans*), and flies (*Drosophila melanogaster*) are viable with only mild phenotypes (Nicolet and Craig, 1989; Song et al., 2009; Ambegaokar and Jackson, 2011). In contrast, the deletion of *STIP1* is lethal early in embryonic development in the mouse (Beraldo et al., 2013), possibly indicating that the function of Hop might be cell type-specific or dependent on the specific cellular state or requirements. In this study, we have explored why Hop is present in eukaryotes, what its critical functions are, and whether and how the eukaryotic Hsp70-Hsp90 molecular chaperone machines may function without Hop to ensure proteostasis. Our studies on the functions of Hop as a co-chaperone of the Hsp70/Hsp90 molecular chaperone machines and facilitator of protein folding and assembly led us to the discovery of alternative cellular strategies that ensure proper protein folding and proteostasis in human and yeast cells lacking this co-chaperone. These findings highlight the persistence of evolutionarily more ancient mechanisms in eukaryotic cells that may contribute to balance protein folding and degradation under certain conditions.

## RESULTS

### Human Hop Knock-out Cells Maintain Cellular Fitness and Proteostasis and Are Not Hypersensitive to Proteotoxic Stress

To study the functions of Hop in eukaryotic cells, we knocked out the gene *STIP1*, which encodes mammalian Hop, in several different human cell lines by using the CRISPR/Cas9 gene editing technique. The absence of the full-length Hop in knock-out (KO) clones was confirmed by immunoblotting using a specific antibody to Hop (Figure 1A). We did note that the HEK293T clone KO1 expresses a residual low level of a truncated form of Hop, which we characterized by mass spectrometry (MS) (Table S1); however, in subsequent experiments presented below, KO1 proved to behave essentially like the other HEK293T clone (KO33), which is devoid of any detectable trace of Hop. The frequency of obtaining KO clones with these human cell lines ranged between 33-46% indicating that Hop is not absolutely essential in human cells similarly to what had previously been found for the budding yeast *Saccharomyces cerevisiae* (Nicolet and Craig, 1989). Morphological examination revealed no obvious differences between wild-type (WT) and KO cells (Figure 1B). Growth rates of KO cells are only moderately reduced compared to WT cells (Figure 1C); this observation was supported by cell cycle analyses in that KO cells show less cyclic phase cells (S - G2/M) and more cells in the G0/G1 resting phase (Figure 1D, Figure S1A-B). The flow cytometric analyses of cells stained with annexin V and propidium iodide (PI) showed that none of the KO cell lines have any obvious viability issues (Figure 1E, Figure S1C-D).

**Figure 1.**
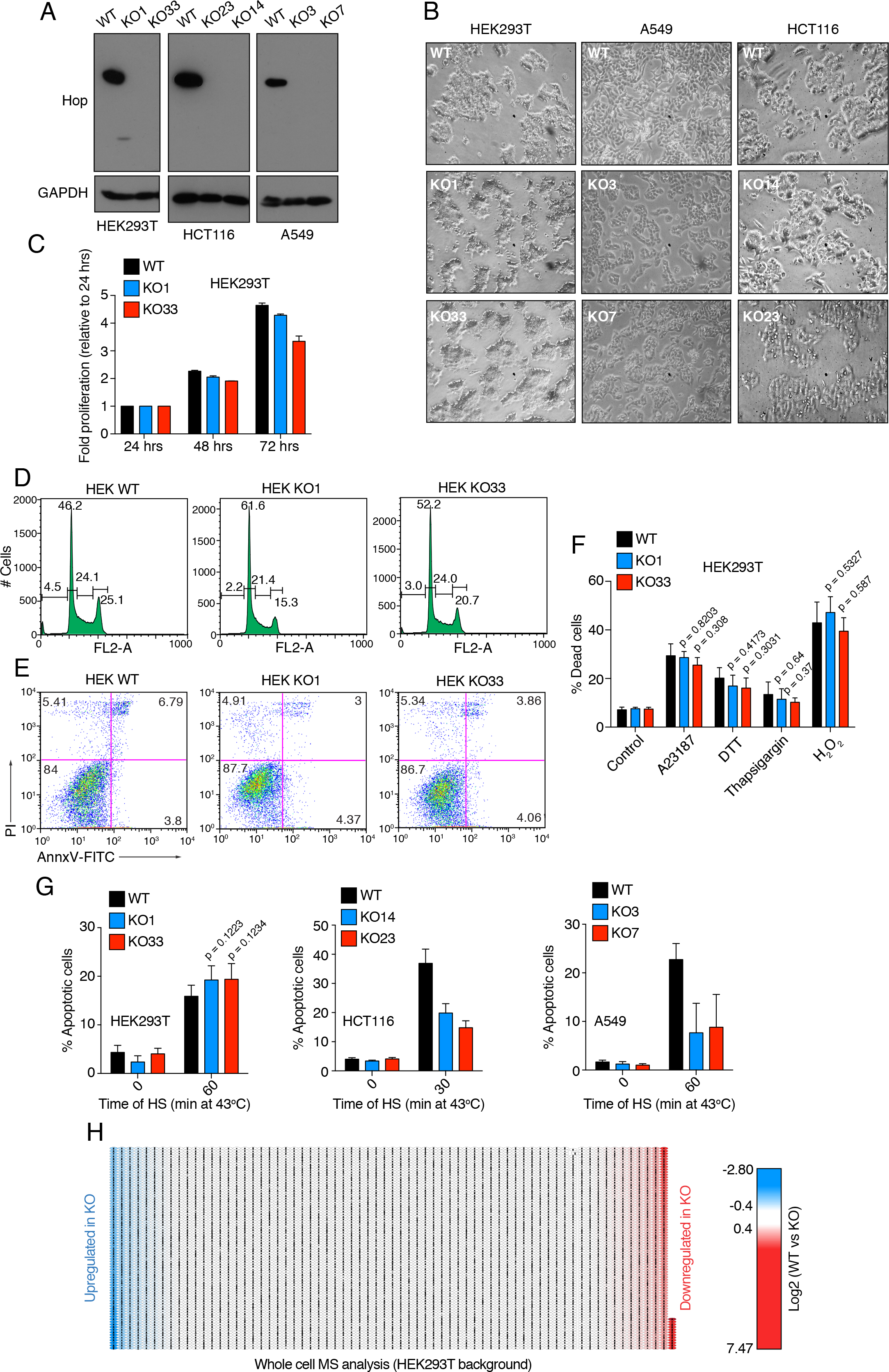
Analyses of Proliferation, Stress Sensitivity and Whole Cell Proteome of Human Hop KO Cells. (A) Immunoblot analysis of two independent KO clones for each cell line. (B) Phase contrast micrographs of WT and KO cells. (C) Cell proliferation analysis of WT and KO cells by MTT assay. (D) Flow cytometry histograms representing cell cycle distributions of WT and KO cells; HEK, HEK293T. (E) Dot plots represent live/dead cell distributions visualized by annexin V-FITC (AnnxV-FITC) and PI staining using flow cytometry. The number in each quadrant gives the % of the total analyzed cell population for a particular sample. (F) Flow cytometric quantification of cell death induced by A23187 (20 *μ*M, 24 hrs), DTT (1 mM, 6 hrs), thapsigargin (1 *μ*M, 24 hrs), and H_2_O_2_ (1 mM, 6 hrs). (G) Flow cytometric quantification of apoptotic cells (sub-diploid cells in cell cycle analysis) after 48 hrs of recovery at 37°C after HS for 0 to 60 min with WT and KO cells. (H) Heat map of whole cell proteome changes. Proteins upregulated and downregulated in KO33 cells are represented in blue (<-0.4 log2 fold changes) and red (> 0.4 log2 fold changes), respectively. Scale bar represents log2 fold changes of the label-free quantification value (LFQ).

We then determined how these KO cells survive in various stress conditions. We exposed WT and KO cells to thapsigargin, DTT, and A23187, which all induce the unfolded protein response of the endoplasmic reticulum, and to the oxidative stress inducer H_2_O_2_. Remarkably, KO cells are clearly not hypersensitive to these stress agents since these lead to comparable numbers of dead cells (Figure 1F) and to a rather reduced impact on cell morphology compared to WT cells (Figure S1E). Upon exposure to heat shock (HS), a stress which induces protein misfolding and aggregation, KO cells showed a similar sensitivity in the HEK293T background, but a markedly reduced sensitivity in the HCT116 and A549 backgrounds (Figure 1G). We also challenged both WT and KO cells with azetidine-2-carboxylic acid (AZC), a toxic analog of proline; incorporation of this proline analog into nascent proteins causes misfolding and subsequent protein aggregation. KO cells are more resistant to this proteotoxic stress, based on the comparison of the cellular morphology of AZC-treated WT and KO cells (Figure S1F).

A label-free MS quantitation of the whole cell proteome indicated that the vast majority of the proteome is unaltered between WT and KO cells (Figure 1H, Figure S1G). With our cut-offs for biologically significant changes of protein levels (see STAR Methods), only those of the top and bottom 4 and 12% of the identified proteins of HCT116 and HEK293T cells, respectively, are altered by the KO. These results indicated that the absence of Hop does not compromise the overall proteostatic equilibrium; instead, if anything, proteostatic buffering of KO cells becomes more robust and resilient to proteotoxic stress conditions.

### The Proteasome and the Hsp70-Hsp90 Molecular Chaperone Axis Are Differentially Required in KO Cells

Cells experience intrinsic or environmental stress-induced proteotoxicity and they notably cope with these stresses by using two important protein quality control mechanisms: degradation of misfolded proteins by the proteasome and in some circumstances by autophagy, and refolding and stabilization of misfolded proteins by molecular chaperones, including the Hsp70-Hsp90 molecular chaperone machines (Figure 2A). To assess how these mechanisms are operating in KO cells, we pharmacologically blocked the functions of Hsp90, Hsp70, the proteolytic activity of the proteasome, and ubiquitination. We observed that KO cells are hypersensitive to Hsp90 inhibition with geldanamycin (GA), as demonstrated by an enhanced cell cycle arrest in the G2/M phase, and enhanced accumulation of apoptotic cells (Figure 2B, Figure S2A). This observation was further confirmed by a striking GA-induced morphological collapse (Figure S2B) and increased accumulation of dead cells with KO cells (Figure S2C). KO cells also showed a higher sensitivity to the chemically different Hsp90 inhibitor PU-H71 (Figure S2D) and the C-terminal Hsp90 inhibitor novobiocin (Figure S2E). Remarkably, these findings are reminiscent of the observation with yeast that Hsp90 inhibition and *Δsti1* are synthetically lethal (Liu et al., 1999), indicating that the synthetic lethality of a *STIP1/STI1* deletion and Hsp90 inhibition is evolutionarily conserved. Similarly to Hsp90 inhibitors, the Hsp70 inhibitor JG-98 blocks the growth of KO cells more efficiently than that of WT cells (Figure 2C, Figure S2F). We conclude from these experiments that both Hsp90 and Hsp70 continue to be functionally required in KO cells, which appear to be even more dependent on these molecular chaperones.

**Figure 2.**
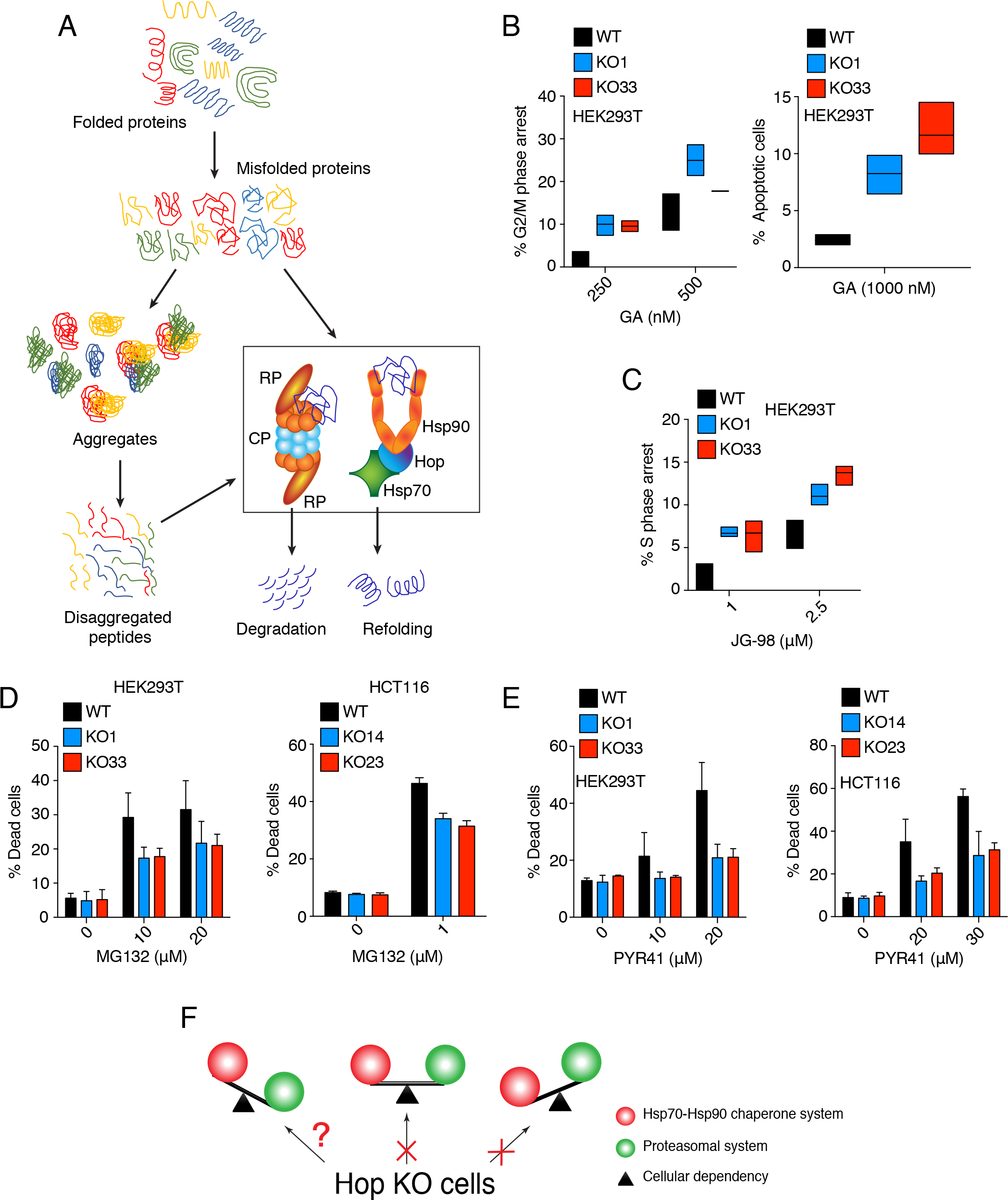
KO Cells Are Differentially Dependent on Proteasome and Hsp70-Hsp90 Molecular Chaperone Functions. (A) Schematic representation of the proteostatic equilibrium relevant to this study. (B) Flow cytometric analysis of the GA-induced G2/M phase cell cycle arrest (left) and apoptosis (right) after 24 hrs of treatment at the indicated concentrations. Data are represented as a box plot. (C) Flow cytometric analysis of the JG-98-induced S phase cell cycle arrest after 24 hrs of treatment at indicated concentrations. Data are represented as a box plot. (D, E) Flow cytometric analysis of the MG132 and PYR41-induced cell death after 24 and 48 hrs of treatment, respectively, of WT and KO cells at the indicated concentrations. (F) Schematic representation of the altered proteostatic equilibrium of KO cells.

To check the other side of the coin of proteostasis, we next performed the cytotoxicity assays with proteasomal inhibitors. The treatments with both MG132 and bortezomib resulted in reduced cytotoxicity with all KO cell lines (Figure 2D, Figure S2G-H) and a less severe impact on cellular morphology than with WT cells (Figure S2I). KO cells also displayed a higher resistance to the E1 ubiquitin ligase inhibitor PYR41 (Figure 2E, Figure S2J). These results suggested that KO cells are less dependent both on ubiquitination and proteasomal degradation. Moreover, transient overexpression of full-length WT Hop in KO cells completely reversed the sensitivity to Hsp90 and proteasomal inhibitors (Figure S2K-M). These results led us to hypothesize that proteostasis in KO cells is ensured by an alternative equilibrium between protein degradation by the proteasome and protein stability/refolding supported by the Hsp70-Hsp90 molecular chaperone machines (Figure 2F). This raises two questions: (1) Is the function of the ubiquitin-proteasome system compromised in the absence of Hop? (2) How can the Hsp70-Hsp90 molecular chaperone machines function efficiently in the absence of Hop?

### The Hsp70-Hop-Hsp90 Ternary Complex Is Physically and Functionally Associated With the Proteasome

To determine whether Hop is responsible by itself for optimal proteasomal function or whether it requires an association with Hsp70 and Hsp90, we carried out immunoprecipitation (IP)-MS experiments with Hop mutants that are defective for Hsp70/Hsp90 binding. Briefly, we generated the TPR1 mutant K8A, which does not bind Hsp70, the TPR2A mutant K229A, which does not bind Hsp90, and the corresponding double mutant (Bhattacharya et al., 2018). We confirmed the expected interaction patterns of these HA-tagged Hop mutants with Hsp70 and Hsp90 by IP (Figure 3A). Based on this result, we defined the Hop-specific interactome by an IP-MS analysis with transfected HEK293T KO cells. After an initial quality control by SDS-PAGE (Figure S3A), samples were subjected to label-free LC/MS-MS analysis. By comparison with the proteins associated with the TPR double mutant, ~41% were identified only with WT Hop and ~57% were enriched with WT Hop (Figure 3B). Thus, the Hop interactome is largely dependent on the ability of Hop to bind Hsp70 and Hsp90. We took a closer look at the Hop interactome by focusing on the components of the Hsp70-Hsp90 molecular chaperone machines. All of these proteins are either enriched or only present with WT Hop, including Hsp70 (HSPA1), Hsc70 (HSPA8), Hsp90α (HSP90AA1), and Hsp90β (HSP90AB1) (Figure 3C). By comparison, Grp94 (HSP90B1), the endoplasmic reticulum-specific Hsp90 isoform, and not a known interactor of Hop, yields the lowest enrichment (~1.5 fold, Figure 3C) with WT Hop in this subset of proteins. When we performed a gene ontology (GO) term enrichment analysis with all preferred interactors of WT Hop using the Enricher web server (http://amp.pharm.mssm.edu/Enrichr), focusing on KEGG and WIKI pathway-annotated proteins, we observed that proteasomal and proteasome-associated ubiquitin-related proteins are overrepresented with WT Hop (Figure S3B-D). Upon searching for all the proteasomal core components and ubiquitin-related proteins, we found that all of these proteins are either enriched or only present with WT Hop (Figure 3D). Interestingly, out of the 19 identified proteasomal components, 16 are associated with the RP of the proteasome (Figure 3D). We reevaluated a published MS dataset of IPs of a panel of proteasomal components from yeast (Guerrero et al., 2008). Remarkably, three of the bait proteins jointly pulled down Sti1 (yeast Hop), Hsp82/Hsc82 (yeast Hsp90s), and Ssa1/2 (yeast Hsp70s), compatible with the conclusion that the ternary complex associates with the proteasome in yeast (Figure S3E). Thus, the Hsp70-Hop-Hsp90 ternary complex is physically associated with proteasomal components, most notably with RP proteins.

**Figure 3.**
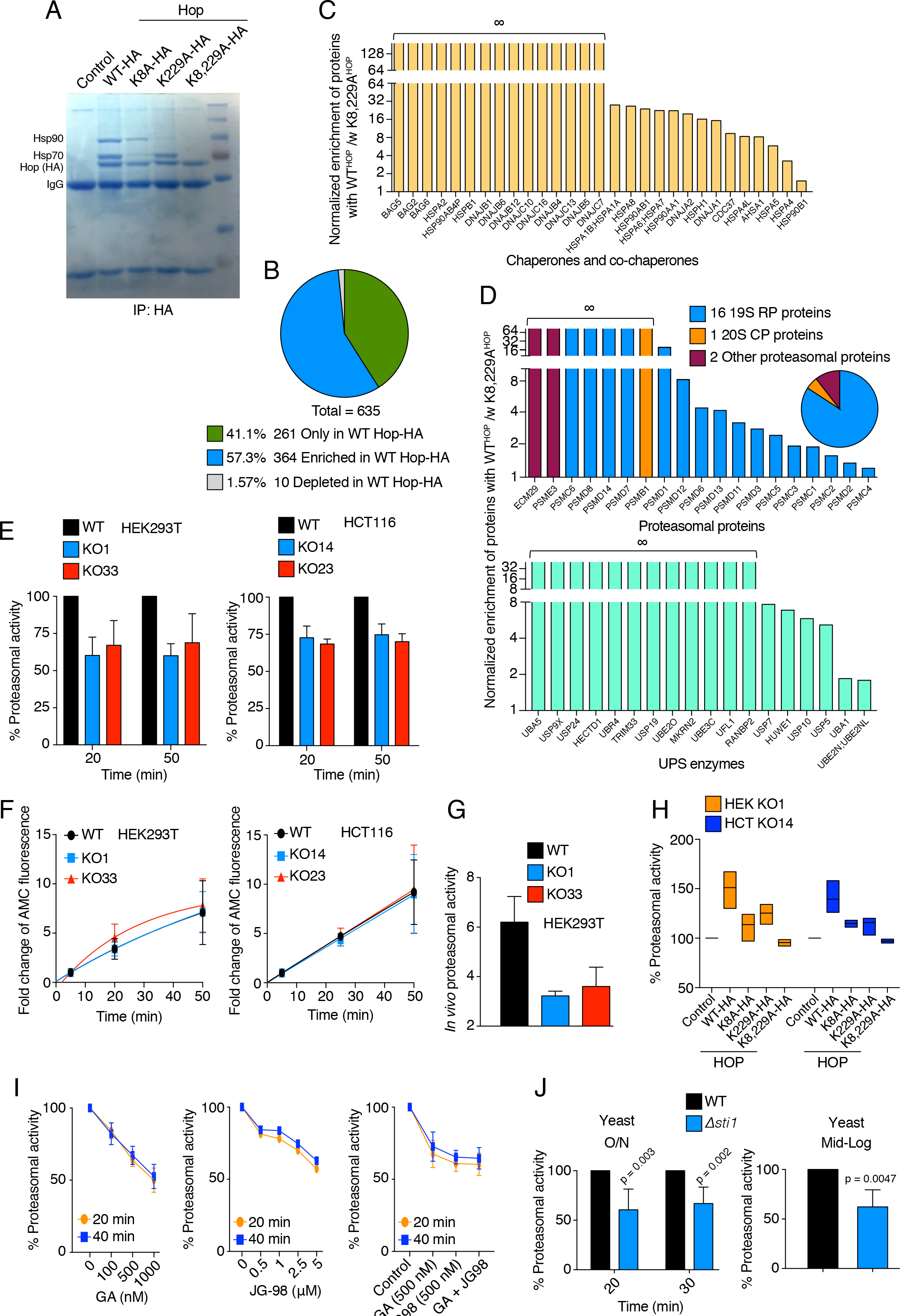
The Hsp70-Hop-Hsp90 Ternary Complex Is Required for Proteasomal Activity. (A) Immunoprecipitation of HA-tagged WT and TPR mutants (K8A, K229A, and K8,229A) of Hop overexpressed in HEK293T KO1 cells with an anti-HA antibody, visualized by Coomassie blue staining of the SDS-PAGE. (B) The pie chart represents the indicated groups of interactors of Hop identified by IP-MS analysis of HA-tagged WT Hop and its TPR double mutant exogenously overexpressed in KO1 cells. (C) The bar graph represents the normalized linear fold changes of Hsp70/Hsp90-related chaperone and co-chaperone proteins immunoprecipitating with the HA-tagged WT Hop compared to its TPR double mutant from KO1 cells by MS analysis as demonstrated in Figure S3A. ∞ signifies proteins only identified with WT Hop. (D) Bars graphs of MS analyses as in panel C highlighting proteasomal proteins (top) and UPS-related enzymes (bottom). Inset: Classification of the identified proteasomal proteins. ∞ signifies proteins only identified with WT Hop. (E) *In vitro* steady state proteasomal activity with extracts of WT and KO cells at the indicated time points after initiation of the reaction with the activity reporter suc-LLVY-AMC. The proteasomal activity of WT cells was considered as 100% for each time point. (F) Rate of proteasomal activity determined with the activity reporter suc-LLVY-AMC. The AMC fluorescence at 5 min was considered as the base value of the reaction, and all other data points for a particular sample were normalized to that. (G) Flow cytometric determination of the *in vivo* proteasomal activity using the Ub-M-GFP and Ub-R-GFP reporter plasmids. *In vivo* proteasomal activity (in arbitrary units.) = % Ub-M-GFP positive cells - % Ub-R-GFP positive cells. (H) *In vitro* steady-state proteasomal activity of extracts from KO cells overexpressing WT Hop or Hop TPR mutants. The proteasomal activity of mock transfected KO cells was considered as 100%. Data are represented as a box plot. (I) Impact of GA (left) and JG-98 (middle, n=2) alone, and the combination of these two inhibitors (right) on the proteasomal activity of extracts from WT cells treated for 24 hrs with the indicated concentrations. Proteasomal activity was measured at 20 min and 40 min after the initiation of the reaction with suc-LLVY-AMC. The proteasomal activity of untreated controls was considered as 100%. (J) *In vitro* steady-state proteasomal activity of extracts of overnight (O/N, left) and mid-log phase (right) cultures of *STI1* (WT) and *Δsti1* yeast cells (BY4741 strain background). The proteasomal activity of WT cells was considered as 100% for each time point. HEK and HCT, HEK293T and HCT116, respectively.

The next experiments were designed to test whether the Hsp70-Hop-Hsp90 ternary complex is functionally relevant for the proteasome. We performed an *in vitro* proteasomal activity assay with extracts from WT and KO cells and noticed that the total steady-state activity of the 26S/30S proteasome is reduced across all KO cell lines (Figure 3E, Figure S3F). In contrast, the rate of proteasomal activity is not significantly altered in KO cells (Figure 3F, Figure S3G). These results suggested that functional 26S/30S proteasome particles might be less abundant in KO rather than less functional on a per particle basis. We confirmed these *in vitro* results with an *in vivo* proteasomal activity assay (Dantuma et al., 2000); this involved the transient expression and flow cytometric quantitation of a degradation-prone ubiquitin-GFP fusion protein (Ub-R-GFP) in parallel to its stable counterpart (Ub-M-GFP) as a control (Figure 3G, Figure S3H). Upon transient expression of WT and TPR mutant Hop in KO cells, the proteasomal activity is rescued by WT Hop, whereas single TPR mutants are not as efficient and the TPR double mutant behaves like a completely dead mutant regarding this function (Figure 3H, Figure S3I). These results established that the Hsp70-Hop-Hsp90 ternary complex is not only physically associated with proteasomal components, but also functionally required for proteasomal activity.

Previous studies in different models had indicated that reduction or deletion of Hsp90 could negatively influence the proteasomal activity (Imai et al., 2003; Yamano et al., 2008; Nanduri et al., 2015; Choutka et al., 2017). When we deleted the genes encoding Hsp90α (*HSP90AA1*) or Hsp90β (*HSP90AB1*) in HEK293T cells, we found a reduced level of steady-state proteasomal activity (Figure S3J). Moreover, the pharmacological inhibition of Hsp70 (with JG-98) or Hsp90 (with GA) in WT cells similarly led to a reduction of proteasomal activity (Figure 3I), whereas proteasomal activity remained unaffected when GA is added to the extracts after cell lysis (Figure S3K). Interestingly, a combination of suboptimal concentrations of JG-98 and GA did not show any additive or synergistic effects in the reduction of proteasomal activity (Figure 3I, right panel). These results collectively indicate that functional Hsp70 and Hsp90, including the possibility to form a ternary Hsp70-Hop-Hsp90 complex, are essential for optimal proteasomal function and activity. Furthermore, we found that the requirement for the Hsp90-Hop-Hsp70 ternary complex for full proteasomal activity is conserved in yeast; analogously to mammalian cells, the steady-state proteasomal activity, but much less so the rate of proteasomal activity, is reduced in a *Δsti1* strain (Figure 3J, Figure S3L).

### Stability of Individual Proteasomal Components Is Independent of the Hsp70-Hop-Hsp90 Ternary Complex

We considered the following two possibilities to explain the impact of the *STIP1(Hop)* KO on the proteasome: (1) Proteasomal components are clients of the ternary complex, and in the absence of Hop, they become unstable and subsequently degraded; (2) the ternary complex is required for 26S/30S proteasome assembly and/or maintenance. To address the first possibility, we reanalyzed previously published datasets of whole-cell proteomic experiments, where HeLa and Jurkat cells had been treated with Hsp90 inhibitors (Sharma et al., 2012; Fierro-Monti et al., 2013). As expected, the levels of some well-known intracellular clients of Hsp90 are decreased and Hsp90-associated molecular chaperone and co-chaperone proteins are increased by Hsp90 inhibition (Figure 4A). In contrast to what we observed with clients, Hsp90 inhibitors do not reduce the protein levels of any of the known core proteasomal proteins (Figure 4A, right panel). We also experimentally checked the impact of GA on the levels of a few RP components in WT cells by immunoblotting with specific antibodies and found that none of them are affected (Figure 4B).

**Figure 4.**
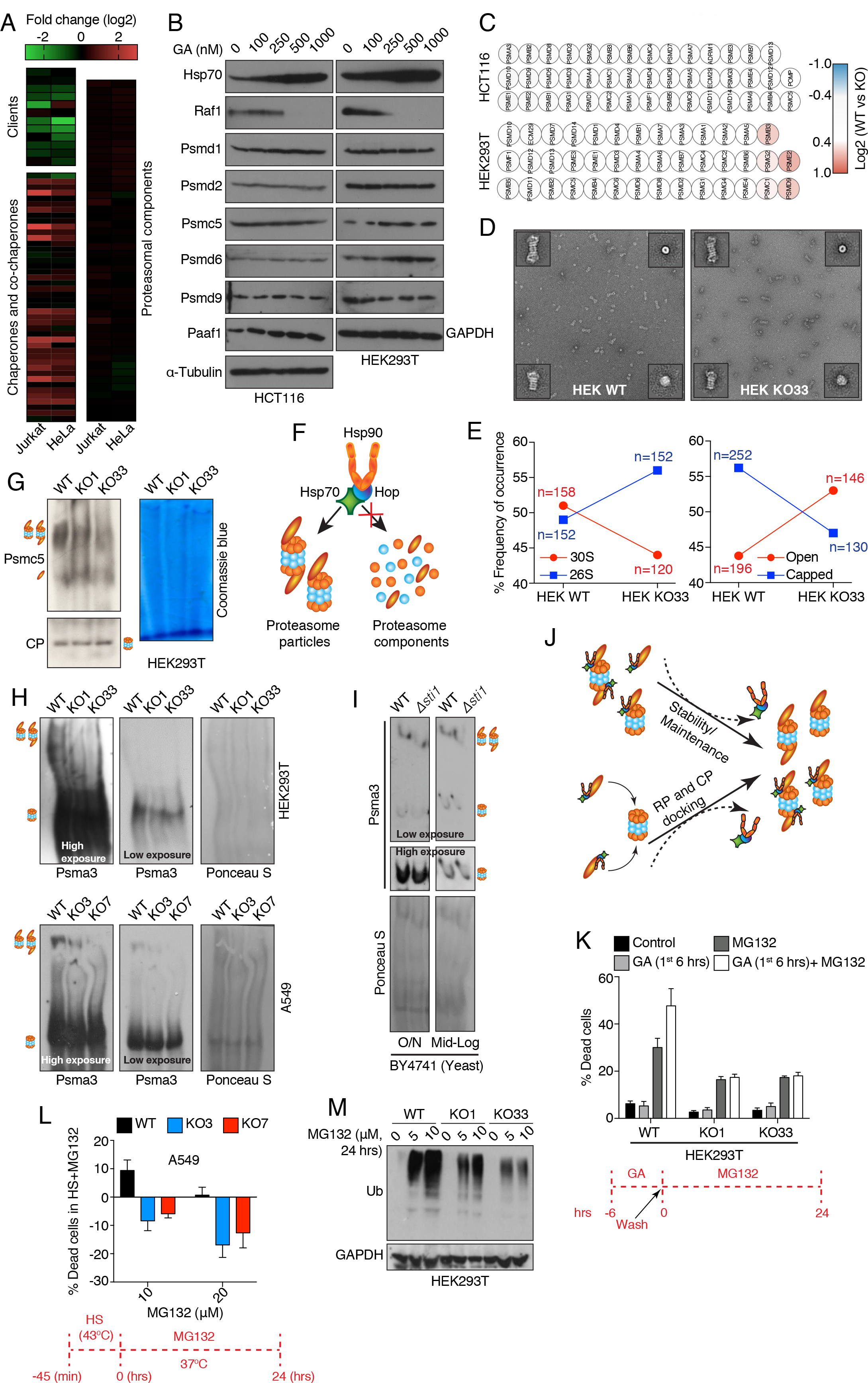
Hsp70-Hop-Hsp90 Ternary Chaperone Complex Is Required for Proteasome Assembly. (A) Heat maps of normalized fold changes of the levels of Hsp90 clients and co-chaperones, and proteasomal components in Jurkat and HeLa cells treated with Hsp90 inhibitors. The MS data are from previous publications (Sharma et al., 2012; Fierro-Monti et al., 2013). Overexpressed and downregulated proteins in the treated datasets are represented in red and green, respectively. (B) Immunoblots of indicated proteasomal subunits from WT cells treated with GA for 24 hrs at indicated concentrations. (C) Heat maps of the normalized fold changes of the levels of proteasomal proteins identified by whole cell MS analyses with WT vs. Hop KO cells. The scale bar represents the log2 fold changes (WT vs KO) of the LFQ values. (D) Images of purified proteasome particles from WT and KO HEK293T cells obtained by negative staining TEM. Insets of each micrograph: side views of double-capped 30S proteasome (top left) and single-capped 26S proteasome (bottom left) particles, and top views of proteasome particles with the CP face up (top right) and the RP face up (bottom right). (E) Frequencies of side (left) and top views (right) of proteasomal structural projections from WT and KO33 HEK293T cells. n represents the total numbers of structural projections of proteasome particles analyzed by TEM; the colored lines only help to visualize the changes. (F) Schematic representation of the involvement of the Hsp70-Hop-Hsp90 ternary complex in proteasome assembly rather than for individual proteasomal proteins. (G-I) Abundance of different proteasomal particles of human and yeast cells as indicated displayed by 4% native-PAGE and subsequent immunoblotting (with antibodies against the indicated proteasome components). Slowly and fast migrating bands represent complexed 26S/30S proteasome particles and the free forms of either the 19S RP (panel G) or 20S CP (panels H, I), respectively. Nitrocellulose filters stained with Ponceau S or Coomassie blue-stained gels indicate equal loading of proteins. (J)Models of how the Hsp70-Hop-Hsp90 ternary complex could enhance the abundance and stability of assembled 26S/30S proteasome particles. (K)Flow cytometric quantification of cell death of WT and KO cells treated with MG132 (10 μM) and GA (500 nM) as indicated in the scheme in red. (L)Flow cytometric quantification of cell death of WT and KO cells treated with MG132 for 24 hrs at the indicated concentrations during the recovery phase at 37°C after a HS (43°C, 45 min). (M)Immunoblot of ubiquitinated-proteins from WT and KO cells treated with MG132 for 24 hrs at the indicated concentrations.

Similarly, we did not find any striking and cell line-independent differences in protein levels of proteasomal proteins between WT and KO cells in the datasets of our own whole-cell proteomic analysis (Figure 4C). Thus, it seems unlikely that proteasomal components are dependent on Hsp90 for accumulation and/or stability.

### Overall Composition of the Assembled Proteasome Is Similar in the KO Cells

To evaluate the impact of the absence of the ternary complex on proteasome assembly/maintenance, we purified 26S/30S proteasome particles from both WT and KO cells. 26S/30S proteasome particles were pulled out by an affinity purification strategy targeting the RP protein S5a (human gene name *PSMD4*); note that this purification scheme enriches for single-(26S) and double-capped (30S) CPs and that any subsequent analysis with this material could not report on free CPs or unassembled proteasome proteins. The integrity of the purified 26S/30S proteasomal complex was analyzed by native gel electrophoresis (Figure S4A-B), and by functional assays without and with MG132 (Figure S4C-D). To characterize the composition of the proteasome purified from WT and KO cells, we performed a comparative label-free LC/MS-MS analysis. We did not find any consistent cell line-independent differences of any identified stoichiometric components of the proteasome between WT and KO cells (Figure S4E-F). Although we noticed that three substoichiometric components and chaperones of the proteasome were reduced in preparations of proteasome particles from HEK293T KO cells (Figure S4F), we did not further consider them since they were unchanged in the HCT116 background (Figure S4E). The aforementioned Hop IP-MS analysis showed that many proteasomal proteins are associated with WT Hop (see Figure 3D). Interestingly, in our own proteomic analyses of purified proteasome from WT HEK293T and HCT116 cells, all components of the Hsp70-Hop-Hsp90 ternary molecular chaperone complex could be identified, albeit at very low substoichiometric levels. This again demonstrates that the Hsp70-Hop-Hsp90 ternary complex is specifically associated with the proteasome, while the low stoichiometry suggests that the association may be regulatory and transient. We concluded from these experiments that overall the composition of fully assembled proteasome particles is similar in the absence of Hop. However, we cannot absolutely rule out the possibility that a minor fraction of purified free RP influenced the overall proteasomal composition.

### Optimal Proteasomal Assembly Requires the Hsp70-Hop-Hsp90 Ternary Complex

To check the impact of the absence of Hop and of the ternary molecular chaperone complex on the structure of the proteasome, we studied the structural integrity of the purified proteasome particles by negative staining transmission electron microscopy (TEM). 2D class averaging of all visible TEM structures led to four different proteasomal structural projections. We could see side views of single-capped 26S and double-capped 30S proteasome particles (Figure 4D, Figure S4G). Moreover, we could observe the two expected top views: an open form of the proteasome with a central hole and a closed form corresponding to CP and RP face up, respectively (Figure 4D). Based on the purification scheme and the relative levels of different forms in the native-PAGE analysis, we did not expect any structures related to free CP (see Figure S4A-B). We measured the dimensions of all four proteasomal projections and confirmed that regardless of the presence of Hop, they are similar to the known values (Walz et al., 1998; Unno et al., 2002) (data not shown). The main difference is the higher abundance of the double-capped proteasome (30S) in proteasome preparations from WT cells while the single-capped proteasome (26S) is more prevalent in preparations from KO cells (Figure 4E, Figure S4H); the opposite situation applies for the relative abundance of open versus capped proteasome particles (Figure 4E). Considering these structural analyses, we hypothesized that Hop, possibly as part of an Hsp70-Hop-Hsp90 ternary complex, is important either for the process of capping of proteasome particles and/or for maintaining the stability of the assembled 26S/30S forms of the proteasome. Collectively we concluded that the individual proteasomal components are not clients of the ternary Hsp70-Hop-Hsp90 chaperone complex, but rather that the ternary complex is involved in the assembly and/or maintenance of the proteasome (Figure 4F).

We next evaluated the ensemble of proteasome particles in whole cell extracts of both WT and KO cells by native-PAGE. We detected RP, CP and 26S/30S proteasome particles using specific antibodies against the CP and RP components Psma3 and Psmc5, respectively, whose total levels are not significantly altered between WT and KO cells (see Figure 4C, Figure S4I). We found that the abundance of the 26S/30S proteasome complexes is reduced in KO cells, whereas free RP and CP are not varying strikingly (Figure 4G-H). We further illustrated this finding by running the native-PAGE with 1.5-fold more total cell extract from KO cells next to an extract from WT cells (Figure S4J). As expected, only WT Hop and not the TPR double mutant can rescue proteasome assembly (Figure S4K-L). Moreover, the reduced proteasome activity in *Δsti1* yeast is mirrored by the reduced assembly and/or stability of proteasome particles (Figure 4I, Figure S4M). Thus, we concluded that the Hsp70-Hop-Hsp90 ternary complex is essential for efficient proteasomal capping and/or for optimal proteasomal stability and maintenance (Figure 4J).

### KO Cells Are Less Dependent on the Proteasome Even With Proteotoxic Stress

Considering the proteasome defect in KO cells, one would expect the degradation flux through the proteasome to be impaired as well. We therefore checked the relative abundance of all identified proteins in the MS datasets of the purified proteasome preparation. This revealed a differential abundance of “proteasomal interactors” between WT and KO samples (Figure S5A). This observation suggested that the utilization and requirement of the proteasome might be different between WT and KO cells. We then filtered the MS data for all proteasomal substrates known to be degraded following either mono- or poly-ubiquitination (Braten et al., 2016). Among all identified mono-ubiquitinated substrates, about 60% are relatively more abundant in proteasome preparations derived from WT cells (Figure S5B). This result indicates that the proteasome is more utilized in WT cells, and thus, that the proteasomal degradation flux might be reduced in KO cells compared to WT cells. However, known poly-ubiquitinated proteasomal substrates are equally abundant in proteasomal preparations of both WT and KO cells (Figure S5C).

So far, we had compared the Hop requirements for the proteasome under normal conditions. We now turned to investigate the effects of proteotoxic stresses. When cells are treated with the Hsp90 inhibitor GA, more proteins form aggregates (Figure S5D), and during a HS, more insoluble and ubiquitinated material accumulates in KO cells (Figure S5E); this demonstrates that the reduction of proteasomal activity and cellular dependency on the proteasome of KO cells is not a symptom of compromised ubiquitination. Upon inhibiting the proteasome in the recovery phase after a 6-hour treatment with GA, we observed that WT cells are dying significantly more than KO cells (Figure 4K, Figure S5F-G). Similarly, inhibiting the proteasome after HS kills WT cells strikingly more than KO cells (Figure 4L, Figure S5H).

To evaluate the proteasomal degradation flux more directly, we blocked the proteasomal activity with MG132 for 24 hrs and looked at the accumulation of ubiquitinated proteins. Significantly less ubiquitinated substrates accumulated in KO cells (Figure 4M), confirming a reduced degradation flux of the proteasome in KO cells since overall ubiquitination is not compromised in the KO cells (see above and Figure S5E). We conclude that WT cells not only have a higher abundance of assembled proteasome particles, but that they are also more dependent on proteasomal function, and even more so in stressed conditions.

### Hsp70 and Hsp90 Functionally Collaborate Without Hop *In Vivo*

Even though the KO cells display a compromised proteasomal assembly and function, overall they nevertheless seem to maintain proteostasis (see Figure 1H, Figure S1G). To provide an additional general assessment of this conjecture, we biochemically fractionated soluble and insoluble proteins from WT and KO cells. Irrespective of genotype, we found similar amounts of ubiquitinated proteins and protein aggregates (Figure 5A; see also untreated samples in Figure S5D-E). Therefore, it seemed conceivable that more efficient chaperoning functions are compensating for the proteasomal defects of KO cells. We started to explore this with *in vivo* luciferase refolding assays; we found a higher rate of refolding of heat-inactivated luciferase in KO cells, consistent with a better chaperoning activity in these cells (Figure 5B, Figure S6A). This higher rate of luciferase refolding in KO cells could be completely reverted to that of WT cells by pharmacological inhibition of Hsp90 or Hsp70 (Figure 5C, Figure S6B). Thus, Hop appears to restrain the refolding activity of Hsp70/Hsp90. Reconstitution experiments with overexpression of WT and TPR mutants of Hop in KO cells showed that only WT Hop inhibits luciferase refolding whereas all TPR mutants do not affect it (Figure 5D).

**Figure 5.**
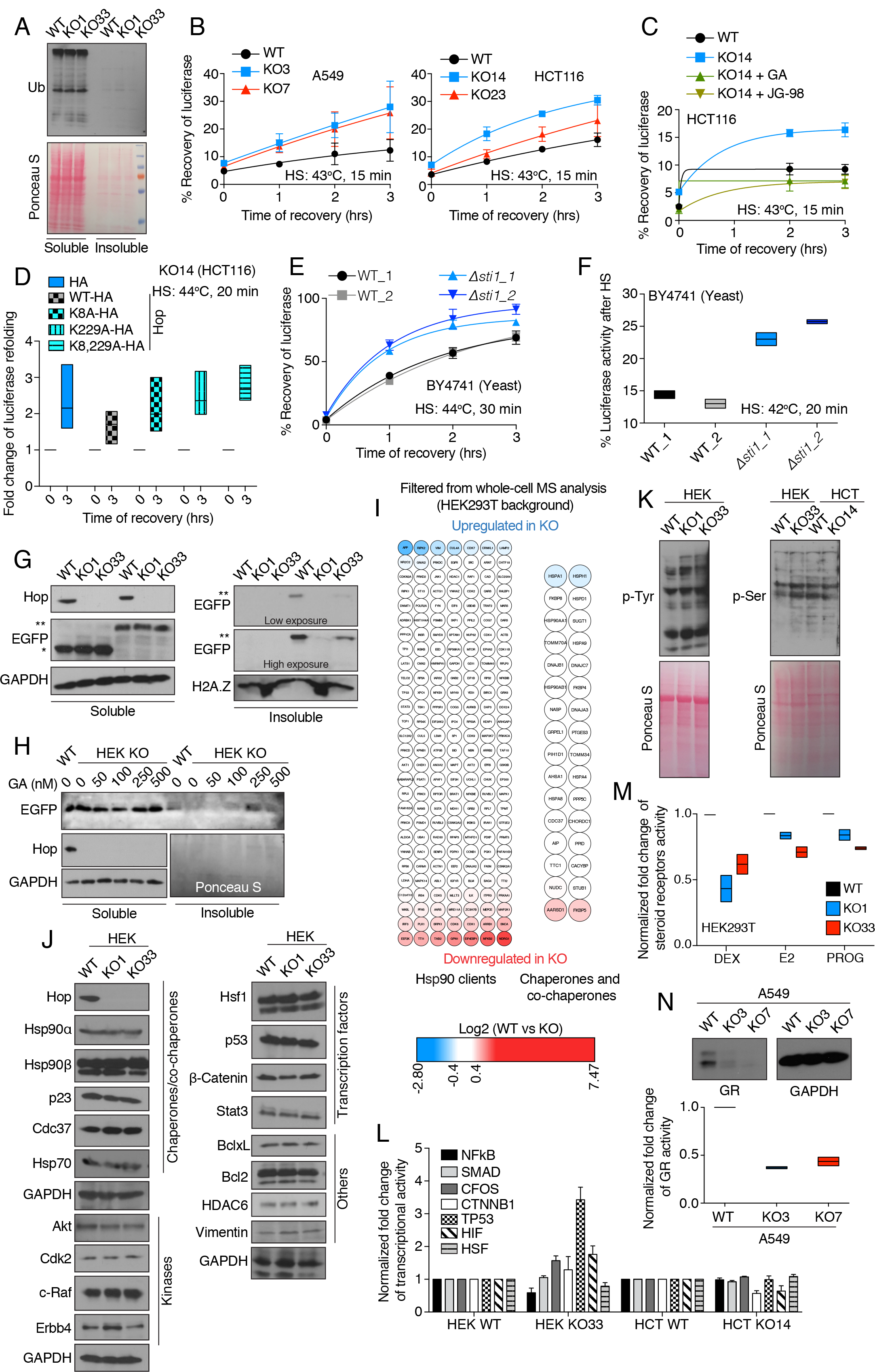
Hsp70 and Hsp90 Are Functional *In Vivo* Even Without Hop. (A) Immunoblot of ubiquitinated proteins from soluble and insoluble protein fractions of WT and KO HEK293T cells. (B) *In vivo* refolding of heat-denatured luciferase in WT and KO cells during the recovery phase at 37°C. The luciferase activity before HS is set to 100%. (C) *In vivo* luciferase refolding in HCT116 KO cells treated with GA (1 *μ*M) or JG-98 (2 *μ*M) during the indicated recovery phase at 37°C. (D) Fold changes of *in vivo* luciferase refolding in HCT116 KO cells exogenously overexpressing WT or TPR mutants of Hop. Non-HS controls for each sample is set to fold 1. Data are represented as a box plot. (E) *In vivo* refolding of heat-denatured luciferase in WT and *Δsti1* yeast cells (BY4741 strain background; two different transformants each) during the indicated recovery phase at 30oC. (F) Residual *in vivo* luciferase activity immediately after mild HS in yeast (n=2). The luciferase activity before HS is set to 100%. Data are represented as a box plot. (G) Solubility of aggregation-prone polyglutamine model protein Q74-EGFP. Immunoblots of Q23-EGFP (first 3 lanes of left and right blots, marked with *) and Q74-EGFP (last 3 lanes of left and right blots, marked with **) from soluble and insoluble protein fractions of WT and KO HEK293T cells. Both overexpressed fusion proteins were detected with an anti-EGFP antibody. GAPDH and H2A.Z serve as loading controls for soluble and insoluble protein fractions, respectively. (H) Immunoblot of Q74-EGFP from soluble and insoluble protein fractions of KO cells treated with GA at the indicated concentrations. The immunoblot of GAPDH serves as loading control for the soluble protein fractions and the Ponceau S staining of the nitrocellulose filter as a loading control for the insoluble protein fractions. (I) Heat maps of the normalized fold changes of the levels of Hsp90 clients (left), and molecular chaperones and co-chaperones (right) identified by whole cell MS analyses. The scale bar represents the log2 fold changes (WT vs KO) of the LFQ values. (J) Impact of the KO on the levels of Hsp90 clients. Immunoblots with antibodies to the indicated proteins present in WT and KO HEK293T cells. (K) Immunoblots of total cellular proteins phosphorylated at either tyrosine or serine residues from WT and KO cells. The nitrocellulose filters stained with Ponceau S indicate equal protein loading. HEK and HCT, HEK293T and HCT116, respectively. (L) Transcriptional activities of the indicated Hsp90 clients determined with corresponding luciferase reporter genes. Data are represented as fold change of transcriptional activities in KO cells in comparison with WT cells (set at 1). NFkB, nuclear factor kappa B; CFOS, c-Fos; CTNNB1, β-catenin; TP53, WT p53; HIF, hypoxia-inducible factor; HSF, heat-shock factor. (M) Transcriptional activities of overexpressed GR (induced by dexamethasone (DEX)), ERα (induced by β-estradiol (E2)), and PR (induced by progesterone (PROG)), as assayed with specific luciferase reporter genes (n=2). Data are represented as fold change relative to those of WT cells (set to 1) as a box plot. (N) Immunoblot of the endogenous levels of GR (top), and its DEX-induced transcriptional activity determined with a corresponding luciferase reporter (bottom, n=2). Data are represented as a box plot. HEK and HCT, HEK293T and HCT116, respectively.

From the literature, *Δsti1* yeast cells are known to be HS-sensitive (Nicolet and Craig, 1989), unlike what we have found for human KO cells (see Figure 1G). In order to check whether the Hop/Sti1-independent chaperoning mechanism is evolutionarily conserved in yeast, we performed an *in vivo* luciferase refolding experiment with the HS-sensitive *Δsti1* yeast strain in the W303 background (Figure S6C). Despite its heat sensitivity, this strain displayed a faster rate of luciferase refolding during the early recovery phase (Figure S6D). To mirror the experiments with our HS-resistant human KO cells, we also performed the experiment with a *Δsti1* mutant yeast strain in the BY4741 background, in which the mutant is as heat-resistant as the WT (Figure S5E). In this genetic background, the enhanced luciferase refolding in the absence of Hop/Sti1 is even more striking (Figure 5E) and there is a much higher residual luciferase activity after a milder HS (Figure 5F). Hence, in the absence of Hop/Sti1, Hsp70 and Hsp90 are functional and responsible for enhanced folding of a model substrate.

Another aspect of chaperoning is keeping aggregation-prone misfolded proteins in a soluble state. Accordingly, we checked the solubility of the aggregation-prone polyglutamine model protein Q74-EGFP (Narain et al., 1999) in WT and KO cells along with a non-aggregating control (Q23-EGFP). We observed a reduction of the aggregated form of Q74-EGFP in KO cells in the HEK293T background (Figure 5G, Figure S6F). This improved solubility seems to be Hsp90-dependent as it could be substantially suppressed by GA (Figure 5H). In HCT116 cells, we did not see any difference in insolubility between the two genotypes (Figure S6G), possibly indicating that cell lines may differ with respect to their intrinsic anti-aggregation activities.

Considering that cell viability and proliferation are relatively unperturbed in the absence of Hop and if indeed Hsp70/Hsp90 function even better for some activities in the absence of Hop, we expected only a minimal impact on Hsp90 clients and co-chaperones. To test this hypothesis, we filtered our whole cell MS dataset for proteins of the Hsp90 interactome (https://www.picard.ch/downloads/Hsp90interactors.pdf) (Echeverria et al., 2011). We observed that the levels of the vast majority of the identifiable Hsp90 interactors are unaffected (Figure 5I-J, Figure S6H). Interestingly, the heat inducible isoform of Hsp70 (encoded by *HSPA1*) and Hsp110 (encoded by *HSPH1*), a nucleotide exchange factor (NEF) of Hsp70, are moderately upregulated in the absence of Hop (Figure 5I, Figure S6H), whereas the expression of other co-chaperones of the Hsp70-Hsp90 chaperone systems, including several J-proteins, remains unaltered (Figure 5I, Figure S6H).

To exclude the formal possibility that Hsp90 clients accumulate to normal levels, but do so in an inactive conformation, we checked the activities of different classes of Hsp90 clients. As there are many tyrosine and serine/threonine kinases amongst the clients of Hsp90, we compared total levels of serine- and tyrosine-phosphorylated proteins between WT and KO cells; we could not see any significant global differences between these two genotypes (Figure 5K). We also did not observe any strong differences between WT and KO cells in the phosphorylation of specific sites indicative of activation of c-Src, an Hsp90 client (Bijlmakers and Marsh, 2000), and of its downstream MAP kinases Erk1/2 (Figure S6I). Using luciferase reporter assays, we checked the transcriptional activities of several transcription factors, which are either Hsp90 clients by themselves or the downstream factor of Hsp90 clients. We found that either the activity is enhanced (in HEK293T cells) or not strikingly compromised (in HCT116 cells) in the absence of Hop (Figure 5L). We conclude that Hsp90 clients not only maintain their steady-state protein levels in KO cells, but that by and large they also maintain their activity compared to WT cells.

### Hop Determines a Unique Spectrum of Hsp90 Client Proteins

Since the vast majority of Hsp90 clients were unaltered in KO cells, we wondered whether challenging their proteostasis by overexpression of Hsp90 clients could reveal Hsp90 vulnerabilities. We overexpressed several steroid receptors, which need Hsp90 for proper folding, stability, and transcriptional activity (Picard et al., 1990; Bohen and Yamamoto, 1993; Nathan and Lindquist, 1995). We observed that the accumulation and transcriptional activity of the glucocorticoid receptor is compromised in KO cells compared to WT cells (Figure 5M, Figure S6J-K). Other steroid receptors, the estrogen receptor α and the progesterone receptor, are only moderately affected in KO cells both in terms of accumulation and transcriptional activity (Figure 5M, Figure S6J, L). The overexpression of v-Src, a viral tyrosine kinase client of Hsp90 (Whitesell et al., 1994), yielded lower protein levels and also strongly reduced kinase activity in KO cells (Figure S6M-N). We overexpressed several other Hsp90 clients (Hif-1α, Hif-2α, and androgen receptor (AR)) and observed that they are not at all compromised in KO cells (Figure S6O). We wondered whether the attenuation of steroid receptor accumulation and activity in KO cells is limited to the case of their exogenous overexpression. At least for GR, this is not the case as we found that both expression and transcriptional activity of endogenous GR are remarkably reduced in A549 KO cells (Figure 5N). Overall, these results suggested that affected steroid receptors, kinases such as v-Src, and the proteasome are rather the exceptions amongst Hsp90 clients with regards to their pronounced Hop-dependence; anecdotally, GR and v-Src are also the very same Hsp90 clients that were the first ones to be discovered and that have been studied for a long time (Picard et al., 1990; Whitesell et al., 1994; Whitesell and Cook, 1996). Moreover, in agreement with a previous publication showing the plasticity of the Hsp90 co-chaperone network for folding of exogenous Hsp90 clients in yeast (Sahasrabudhe et al., 2017), we hypothesized that Hop might determine unique criteria of client selectivity for Hsp90 even in higher eukaryotes.

### An Alternative Prokaryote-like Hsp70-Hsp90 Binary Complex Maintains the Hsp90 Interactome

KO cells maintain proteostasis even though the proteasome function is inefficient; the latter may be counterbalanced by more efficient Hsp70/Hsp90-dependent chaperoning (Figure 6A). Since bacterial Hsp90 and Hsp70 orthologs HtpG and DnaK, respectively, can physically and functionally interact without a Hop-like protein (Genest et al., 2015; Kravats et al., 2017), we investigated the possibility that an alternative Hop-independent Hsp70-Hsp90 molecular chaperone complex might form in human cells. We performed an Hsp90-IP-MS analysis and found Hsp70/Hsc70, encoded by *HSPA1* and *HSPA8*, among the top hits even in KO cells, despite a 4-10-fold reduction (Figure 6B, Figure S7A). We confirmed this Hop-independent interaction of Hsp70 and Hsp90 by a targeted co-IP experiment (Figure 6C, Figure S7B).

**Figure 6.**
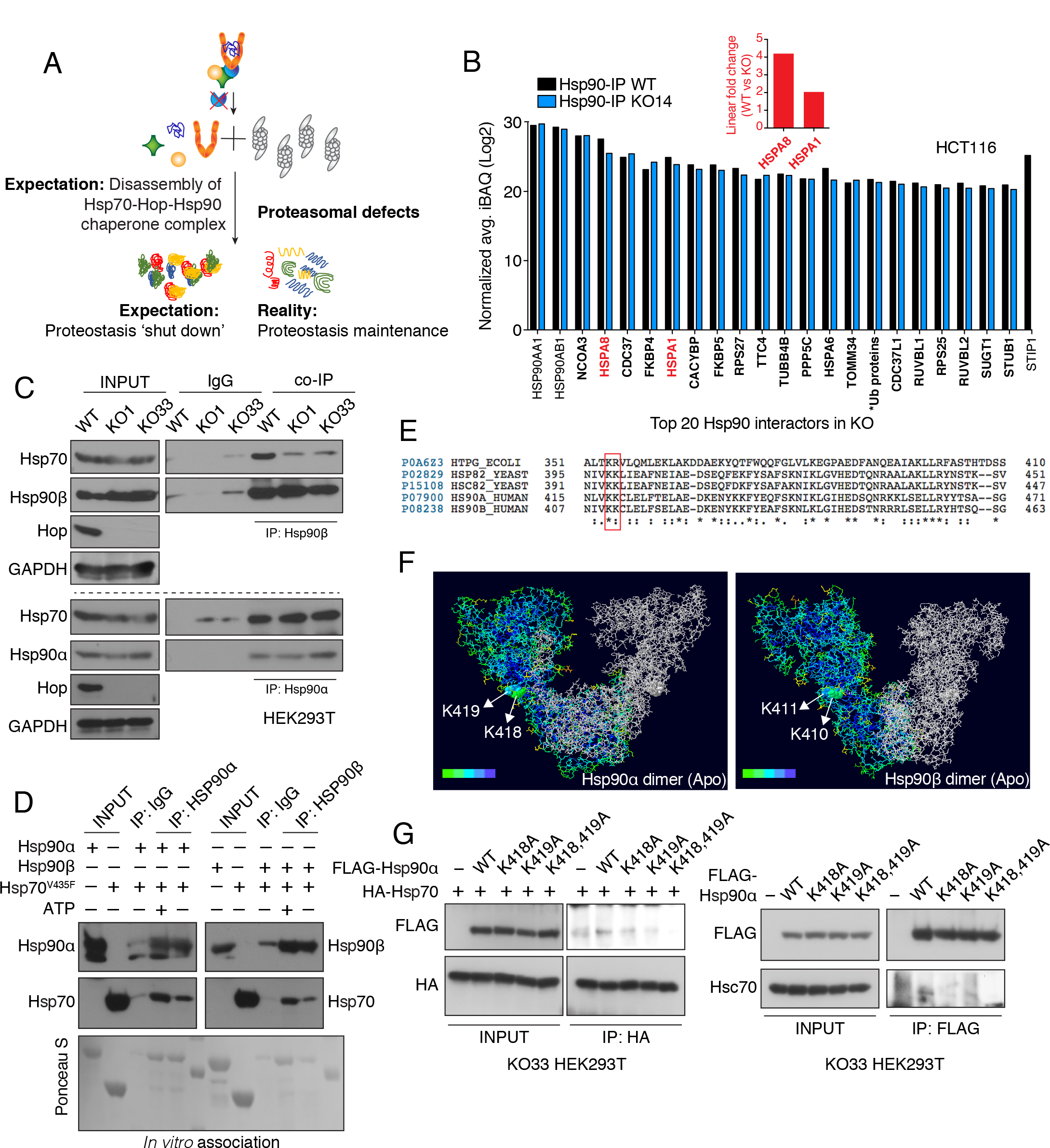
Human Hsp70 and Hsp90 Directly Interact to Form a Prokaryote-like Molecular Chaperone Complex in the Absence of Hop. (A) Model of the impact of removing Hop on the Hsp70-Hsp90 molecular chaperone systems, the proteasome, and proteostasis․. (B) Abundance of the top 20 Hsp90 interactors (those with the highest iBAQ values) of KO HCT116 cells (names in bold). The graph shows the iBAQ values as log2 of the Hsp90-IP-MS analyses. Hsp90α (HSP90AA1) and Hsp90β (HSP90AB1) were the bait proteins, and absence of Hop (STIP1) serves as quality control marker for KO cells. Inset: Linear fold changes of the values for Hsp70 (HSPA1) and Hsc70 (HSPA8). *Ub proteins: UBB, UBC, UBA52, RPS27A. (C) *In vivo* interaction of Hsp90 and Hsp70 as determined by an immunoprecipitation experiment. A control immunoprecipitation was performed in parallel with a normal IgG. (D) *In vitro* interaction of purified recombinant Hsp90 and Hsp70. The substrate-binding mutant V435F of Hsp70 was used in this experiment. (E) Sequence alignments of yeast and human Hsp90 proteins with the bacterial Hsp90 HtpG. Evolutionarily conserved amino acids involved in the direct interaction of HtpG with Hsp70 (DnaK) in bacteria are highlighted by a red box. (F) Surface accessibility of the highlighted amino acids in the predicted dimeric structures (apo form; ADP bound open conformation) of human Hsp90α and Hsp90β. The heat map represents a gradient of accessibility of amino acids in the protein structure with highly accessible ones in green and completely buried ones in dark blue. (G) Interaction of Hsp90 point mutants with Hsp70. Left part: Immunoprecipitation of exogenously expressed HA-tagged WT Hsp70 with coexpressed FLAG-tagged WT or point mutant Hsp90α as indicated. Right part: Immunoprecipitation of exogenously expressed FLAG-tagged WT or point mutant Hsp90α with endogenous Hsc70.

In search of a mechanism promoting a Hop-independent interaction of Hsp70 and Hsp90, we checked whether Hop activity might be functionally redundant in human cells. We revisited our own Hsp90-IP-MS datasets and extracted the data for proteins with a Hop-like architecture, that is with multiple TPR domains. Note that there are only very few of these proteins, which are enriched with Hsp90 in the absence of Hop, and they are very moderately so (Figure S7C). Most importantly, all of these "enriched" TPR proteins are present at very low levels compared to the abundance of Hsp90 and Hsc70/Hsp70 in total cellular extracts from either WT or KO cells, and to Hop in WT cells (Figure S7D). Thus, it is very unlikely that any of these proteins could substitute for Hop for the formation of another Hsp70-(multiple TPR protein)-Hsp90 ternary complex.

These results led us to speculate that human Hsp70 and Hsp90 could directly interact as they do in bacteria (Genest et al., 2015; Kravats et al., 2017) and in *Δsti1* yeast (Kravats et al., 2018). We tested this notion using an *in vitro* association assay with recombinant proteins. To minimize client-type interactions of Hsp70, we used the substrate-binding mutant V435F of Hsp70. The co-IP experiment revealed a direct interaction between Hsp90 and Hsp70 both in the presence and absence of ATP (Figure 6D). We then checked the involvement of evolutionarily conserved surface residues that had been shown to be important for the direct interaction of Hsp70 and Hsp90 orthologs in bacteria (Genest et al., 2015) (Figure 6E). As in bacteria, these residues are well surface-accessible in both human Hsp90 isoforms (Figure 6F, Figure S7E), and modifying these residues in human Hsp90α strongly suppresses the interaction with Hsc70/Hsp70 in KO cells (Figure 6G, Figure S7F).

Our whole-cell MS analysis had already indicated that the binary Hsp90-Hsp70 complex may be sufficient to support the Hsp90 interactome (see Figure 5I, Figure S6H). To address this issue more directly, we compared the levels of all proteins in the Hsp90-IP-MS datasets between WT and KO cells. The majority of the interactions do not change and there are similar proportions of proteins that are enriched or depleted in the absence of Hop (Figures 6B and 7A, Figure S7A and S7G). Thus, client-Hsp90 and co-chaperone-Hsp90 interactions are largely maintained in KO cells, which supports the conclusion that the prokaryote-like Hsp70-Hsp90 binary complex in human cells is functional.

### The Prokaryote-like Human Hsp70-Hsp90 Complex Is Also More Efficient *In Vitro*

Bacterial Hsp70 collaborates with the bacterial J-domain protein Hsp40 and a NEF for the initial substrate recognition, and further substrate folding can be obtained by the collaboration of bacterial Hsp70 with Hsp90 (Mayer and Le Breton, 2015; Moran Luengo et al., 2018). To evaluate the chaperoning activity of the binary human Hsp70-Hsp90 complex, we performed *in vitro* luciferase refolding experiments with purified components. Human Hsp70 was complemented with a J protein, the NEF Apg2, and human Hsp90α. To determine the impact of Hop on this molecular chaperone system, we measured luciferase refolding as a function of increasing concentrations of Hop. To our surprise, but in agreement with our *in vivo* experiments (see Figure 5B, Figure S6A), we discovered that the Hsp70-Hsp90 molecular chaperone systems refold heat-denatured luciferase most efficiently in the absence of Hop (Figure 7B). Increasing concentrations of Hop gradually decrease the final yield of luciferase refolding achieved by the Hsp70-Hsp90 system (Figure 7B-C). In conclusion, human/eukaryotic Hsp70 and Hsp90 can form an evolutionarily conserved chaperone complex that is fully functional for protein folding and maintains the proteostatic equilibrium despite proteasomal assembly defects in the absence of Hop/Sti1.

**Figure 7.**
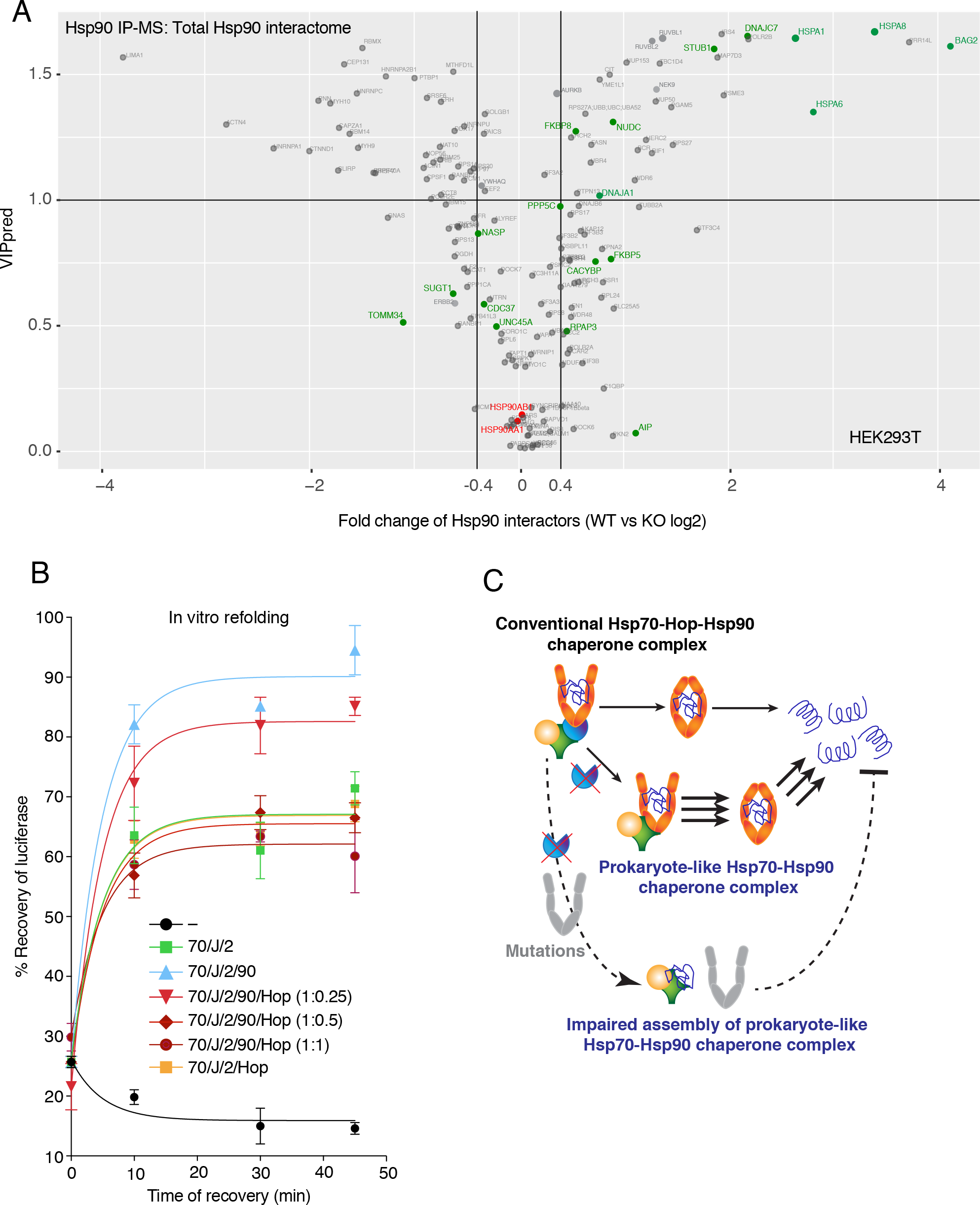
Prokaryote-like Hsp70-Hsp90 Molecular Chaperone Complex Is Functional Both in Human Cells and *In Vitro*. (A) The volcano plot represents the normalized fold changes of the Hsp90 interactors identified by the Hsp90-IP-MS analysis versus their VIP values derived from the OPLS-DA model. Proteins with VIPpred > 1.0 are considered as variables of interest to explain the difference between the conditions (WT vs KO). A value of > 0.4 or < - 0.4 can be considered biologically significant. Hsp90 related co-chaperones are highlighted in green and the two cytosolic Hsp90 isoforms in red. (B) *In vitro* refolding of heat-denatured recombinant luciferase by the indicated combinations of recombinant human Hsp90α (90), Hsp70 (70), DnaJB1 (J) and Apg2 (2) in absence or presence of different concentrations of recombinant Hop. (C) Model comparing the functions of Hsp70-Hsp90 complexes with and without Hop.

## DISCUSSION

Depending on cellular states and stresses, the proteostatic equilibrium can be shifted to alternate mechanisms. For example, HS or inhibition of Hsp90 strongly induce the Hsp70/Hsp40 chaperone system and small Hsps to meet the new cellular requirements (Sittler et al., 2001; Richter et al., 2010; Neckers and Workman, 2012). Here we establish cellular models with human cell lines and yeast, which, in the absence of Hop/Sti1, adopt the more ancient and more efficient mechanism of chaperoning of bacteria and thereby compensate for proteasomal defects, reestablishing an alternate proteostatic equilibrium. Thus, depending on cell-specific requirements or external inputs, eukaryotic cells may still be able to shift to a more prokaryote-like mode of operation for the Hsp70/Hsp90 systems. We speculate that the "invention" of Hop during the evolution from prokaryotes to eukaryotes may have promoted a shift from a proteostatic system centered on refolding to a more extensive use of proteasomal degradation.

Impaired proteasomal function could potentially be compensated by increased autophagy. Proteotoxic stress induced by inhibition of the proteasome can activate autophagy by phosphorylating the autophagy receptor p62 (Lim et al., 2015). The genetic suppression of Hsp90 function in *Drosophila melanogaster* shifts the protein degradation balance from proteasome-mediated degradation to autophagy (Choutka et al., 2017). We cannot formally rule out that increased autophagy might contribute to the new proteostatic equilibrium in Hop KO cells, but we have not seen any upregulation of lysosomal or autophagy-related proteins in our whole cell proteomic data.

Hsp90 itself had already been linked to proteasomal integrity and activity (Imai et al., 2003; Lander et al., 2012). Our data complement this early evidence by demonstrating that it is the Hsp70-Hop-Hsp90 ternary complex that is required for optimal 26S/30S proteasomal assembly and/or for maintaining the proper abundance. According to current models, the RP ATPase-ring docks to the CP α-ring to form the functionally active proteasomal holoenzyme (Murata et al., 2009; Gallastegui and Groll, 2010). We speculate that the ternary molecular chaperone complex primarily binds to the RP, facilitates the docking with the CP and stabilizes the assembled 26S/30S proteasome. The substoichiometric proteasome component Ecm29, which itself is not dependent on Hop (see Figure S4E-F), has been shown to be a tethering factor for RP and CP (Leggett et al., 2002). We propose that the ternary molecular chaperone complex plays a similar role; further studies are required to define more clearly how the ternary complex promotes RP-CP docking and what the interplay with Ecm29 might be. Our MS analysis showed that all components of the Hsp70-Hop-Hsp90 ternary complex are associated with 26S/30S proteasome particles at substoichiometric levels; hence, it is conceivable that the ternary complex acts like other *bona fide* proteasome-dedicated chaperones, which dissociate from the functionally mature proteasomal holoenzyme (Roelofs et al., 2009). Although proteasomal activity and assembly is known to be regulated by posttranslational modifications of its components (Rousseau and Bertolotti, 2018), it is not likely that this can help to explain the proteasomal defects of KO cells; the vast majority of proteins in general and several of the known posttranslational regulators of proteasomal components such as MAPKs, mTORC1-related proteins, and the catalytic subunit of protein kinase A, in particular, are unaltered in KO cells. Thus, we conclude that Hop, as part of the Hsp70-Hop-Hsp90 complex, promotes the assembly of RP with CP and/or contributes to maintaining their assembled state.

Our most surprising finding is that cells lacking Hop/Sti1 compensate the proteasomal defect by improved protein folding. In view of the well-established role of Hop for substrate transfer between the Hsp70 and Hsp90 molecular chaperone machines, and as allosteric regulator of the ATPase activity of Hsp90 (Richter et al., 2003; Alvira et al., 2014; Kirschke et al., 2014; Rohl et al., 2015), this is the last thing one would have expected. Under normal conditions, one can imagine that Hop does promote substrate transfer from Hsp70 to Hsp90, but then ends up stalling the Hsp90 complex, at least temporarily, thereby slowing down further substrate folding. This may be beneficial or even essential to allow the folding or assembly of some substrates such as GR, v-Src, the proteasome, and possibly of some particularly labile clients such as the ΔF508 CFTR mutant (Bagdany et al., 2017). For these, Hop must be able to form the ternary complex. TPR mutants of Hop, which neither bind to Hsp70 nor to Hsp90 nor both are unable to rescue any phenotype in KO cells. This is further supported by the finding that the fortuitously incomplete KO of the HEK293T cell clone KO1 has the exact same phenotype as other clones with a complete KO. The residual ~36 kDa fragment of KO1 only contains the Hsp90-binding domain TPR2A and does not suffice to rescue the KO defects. In contrast, for most Hsp90 clients, the temporary stalling or slow-down of the Hsp90 molecular chaperone machine by Hop may be counterproductive. What defines the Hop requirement of Hsp90 clients remains to be deciphered, but it is clear from our data that the vast majority of Hsp90 clients can be efficiently processed by the prokaryote-like Hsp70-Hsp90 binary complex. We assume that this core molecular chaperone complex still requires the support of other co-chaperones like Hsp40, Cdc37, Aha1 and p23 to select, to process and to release Hsp90 clients (Li et al., 2012; Dunn et al., 2015; Keramisanou et al., 2016; Schopf et al., 2017; Bachman et al., 2018). Compromising the function of these co-chaperones could be particularly detrimental for Hop KO cells running on a somewhat hyperactive Hsp90 machine; indeed, the deletion of *SBA1* (encoding the yeast p23 ortholog) in a Δsti1 yeast strain severely reduces cellular fitness (Fang et al., 1998).

Our data demonstrate that the Hsp70-Hsp90 binary system not only works, but somehow manages to be more efficient. Our quantitative Hsp90-IP-MS experiments show that the steady-state association of Hsp70 and Hsp90 is less prominent. This is compatible with previously reported Kd values for the interactions that are relevant in this context; compared to the Kd for the direct interaction of yeast Hsp70 and Hsp90 of 14 *μ*M (Kravats et al., 2018), the values for the yeast or human pairs Hop-Hsp70 and Hop-Hsp90 are considerably lower (Mayr et al., 2000; Brinker et al., 2002; Wegele et al., 2003). We propose that the binary Hsp70-Hsp90 complex, albeit less stable, is more dynamic than the ternary molecular chaperone complex. Moreover, similarly to what has been reported for the bacterial DnaK-HtpG system (Nakamoto et al., 2014; Genest et al., 2015), the stimulation of the ATPase of one partner by the other might contribute to the improved chaperoning activity in the human system.

Whether Hsp70 and Hsp90 can form the binary molecular chaperone complex in the presence of Hop in WT cells and whether this alternative molecular chaperone complex has any specialized functions in normal or stressed conditions remain open questions. Our data support this hypothesis since we observe a colocalization of Hsp70, Hsp90, Hsp40, and Hsp110, but not Hop, in HS-induced protein aggregates (Figure S7H). However, the specific detection of the Hsp70-Hsp90 binary complex and its characterization in the presence of Hop is technically extremely challenging. Further methodological developments will be necessary to explore the functions of this prokaryote-like complex across different cell types and cellular conditions.

The highly proliferative state of embryonic stem cells is associated with rapid protein turnover and high proteasomal activity (Vilchez et al., 2012). Deletion of some proteasomal subunits has been demonstrated to cause lethality in the mouse, flies and in plants (Sakao et al., 2000; Szlanka et al., 2003; Brukhin et al., 2005; Hamazaki et al., 2007; Beraldo et al., 2013). It is therefore conceivable that the embryonic lethality of the Hop KO in the mouse (Beraldo et al., 2013) could be explained by the failure to meet this particular requirement in embryonic stem cells or in other rapidly proliferating cells during early development. Moreover, the absence of Hop might also compromise the folding/assembly of one or a few specific Hsp90 clients other than the proteasome, which are essential during early embryonic development. What the mouse model clearly demonstrates is that every eukaryotic cell may not be able to shift its proteostatic equilibrium to the overall more efficient chaperoning by the prokaryote-like Hsp70-Hsp90 binary complex in the absence of Hop. A systematic effort will be needed to determine whether Hop KO mouse embryos die because of a proteostasic collapse or because of a more subtle defect relating to some Hsp90 clients. It should be emphasized that the presence of a single *STIP1* allele is sufficient for embryonic development, even though the reduced Hop levels of heterozygote mice does lead to increased cellular stress and vulnerability to ischemia (Beraldo et al., 2013). In this study, we established that the absence of Hop does not affect proteostasis in cellular models. However, cellular or organismic fitness could nevertheless be negatively affected in the long run. Indeed, even though Hop KO worms are viable, their lifespan is reduced (Song et al., 2009). Furthermore, Hop levels drop in the aging brain, and aging is directly correlated with reduced proteasomal and chaperoning activity (Brehme et al., 2014; Rousseau and Bertolotti, 2018). Since the absence of Hop generates a mixed outcome in different experimental models, tissue- and cell type-specific functions of Hop for the maintenance of proteostasis should be further studied.

Our discoveries may also have translational potential. If Hop could be specifically inhibited, this might promote superior chaperoning by the prokaryote-like Hsp70-Hsp90 binary complex. This could be a useful strategy to reequilibrate the proteostatic balance in diseases or altered physiological states with proteasomal dysfunctions. Neurodegenerative disorders such as Huntington’s and Parkinson’s diseases, which are associated with inefficient chaperoning (Labbadia and Morimoto, 2015; Brehme et al., 2014), could potentially benefit from improved chaperoning induced by Hop inhibitors. Some compounds have been reported to inhibit specifically the interaction between Hsp90 and the TPR2A domain of Hop (Yi and Regan, 2008; Pimienta et al., 2011) and may be promising leads for such a therapeutic strategy.

## ACKNOWLEDGMENTS

We are grateful to Alfred L. Goldberg, Matthias P. Mayer, Ueli Schibler, Marcello Maggiolini, Donald McDonnell, Yoshihiko Miyata and Adrienne Edkins for gifts of plasmids. We thank David O. Toft for gifts of antibodies, Jason Gestwicki for the gift of JG-98, and Stacey Mattison for critically reading the methods section of this manuscript. We are also indebted to various previous members of the Picard laboratory for miscellaneous reagents. This work has been supported by the Swiss National Science Foundation and the Canton de Genève.

## AUTHOR CONTRIBUTIONS

K.B. conceived the study, designed and performed experiments, analyzed the data, prepared figures, and wrote the manuscript. L.W. and M.Q. conducted the proteomic analyses. T.M.L and S.G.D.R. designed and performed *in vitro* luciferase refolding assays. P.C.E. performed bioinformatics analyses of proteomic datasets. M.V. generated α- and Hsp90β-KO cells. L.B. contributed to experiments with recombinant proteins. D.W. helped with all yeast experiments. C.B. contributed to TEM experiments and analyses. D.P. conceived the study, contributed to designing the experiments and analyzing the data, supervised the work, and wrote and edited the manuscript. All authors provided critical analysis of the data and contributed to the editing of the manuscript.

## STAR Methods

### CONTACT FOR REAGENT AND RESOURCE SHARING

Further information and requests for resources and reagents should be directed to and will be fulfilled by the Lead Contact Didier Picard.

### EXPERIMENTAL MODELS AND SUBJECT DETAILS

#### Cell lines and cell culture

HEK293T human embryonic kidney cells, HCT116 human colorectal carcinoma cells and A549 human lung epithelial carcinoma cells (as well as the corresponding Hop KO cell lines) were maintained in Dulbecco’s Modified Eagle Media (DMEM) supplemented with GlutaMAX, 10% foetal bovine serum (FBS) and penicillin/streptomycin (100 u/ml) with 5% CO_2_ in a 37 °C humidified incubator. Yeast strain BY4741 (*his3Δ1 leu2Δ0 met15Δ0 ura3Δ0*) and the *Δsti1* variant Y01803 (*his3Δ1 leu2Δ0 met15Δ0 ura3Δ0 Δsti1::kanMX4*), strain W303 (= BMA64-1A) (*ade2-1 can1-100 his3-11,15 leu2-3, 112 trp1-1 ura3-1*) and the *Δsti1* variant W303*Δsti1* (*ade2-1 can1-100 his3-11,15 leu2-3, 112 trp1-1 ura3-1 Δsti1::KanMX4*) were maintained on yeast extract peptone dextrose (YPD) agar plates or in broth at 30°C. To generate the W303 *Δsti1* strain, W303 cells were transformed with a PCR fragment encompassing the *Δsti1::KanMX4* locus of BY4741 and selected with Geneticin (G418, 200 µg/ml) on YPD agar plates. Deletion of *STI1* was verified by PCR amplification and by immunoblot analysis with a specific antibody to the Sti1-TPR2A/B domain. BY4741 and W303 yeast cells were transformed with pLG/LUC, a galactose-inducible luciferase expression plasmid, to produce yeast strains expressing firefly luciferase.

### METHOD DETAILS

#### Plasmids

Site-directed mutagenesis was performed on pcDNA3.1(+)-HOP (WT) to generate human Hop mutants K8A (codon change: AAG > GCG), K229A (AAA > GCA), and Hop mutant K8A/K229A; each with a C-terminal HA-tag (Bhattacharya et al., 2018). Site-directed mutagenesis was performed on the bacterial expression vector pCA528-Hsp90α (WT) and mammalian expression vector pFLAG-CMV2/Hsp90α (WT) to generate the human Hsp90α mutants K418A (codon change: AAA > GCA), K419A (AAA > GCA), and K418A/K419A; each with either a 6x-His tag or FLAG tag. pcDNA3.1(+)-HA-Hsp70 was generated by cloning the human Hsp70 coding sequence, with an N-terminal HA-tag, between *Bam*HI and *Eco*RI restriction sites of the pcDNA3.1(+) vector. Sequences of all oligos (Microsynth) are described in the Key Resources Table.

#### Genome engineering

Hop (*STIP1*)-KO HEK293T, HCT116, and A549 cells; as well as Hsp90α (*HSP90AA1*)/Hsp90β (*HSP90AB1*)-KO HEK293T cells were generated by CRISPR/Cas9 gene editing technology. The guide RNAs (gRNAs) were identified by and the corresponding oligos designed using the ATUM (previously DNA2.0) CRISPR design tool (https://www.atum.bio/). The gRNA sequences corresponding to the 5^th^ exon of *STIP1*, 1^st^ exon of *HSP90AA1,* and 6^th^ exon of *HSP90AB1* are described in the Key Resources Table. Sense and antisense oligos were synthesized (Microsynth), annealed and cloned into the *Bbs*I site of px459 (Addgene plasmid #48139) as described previously (Ran et al., 2013). WT HEK293T, HCT116, and A549 cells were transiently transfected with gRNA containing CRISPR plasmids by polyethylenimine (PEI). At 48 hrs post-transfection, transfected cells were selected with puromycin (3-5 µg/ml). Surviving cells were further cultured in the absence of puromycin until visible cellular foci formed. Cellular foci were individually picked and analyzed by immunoblotting using primary antibodies specific to Hop, Hsp90α, and Hsp90β. Clones which did not express Hop, Hsp90α or Hsp90β were considered the KO cells and frozen in liquid nitrogen. KO clones were further validated by MS analyses to validate the absence of the target protein.

#### Induction of cellular stress and heat shock (HS)

Human cells seeded at a density of 2 × 10^5^ were treated with several intracellular stress inducers to evaluate the effect on cell death and cellular morphology: 1 *μ*M thapsigargin for 24 hrs, 20 *μ*M A23187 for 24 hrs, 1 mM dithiothreitol (DTT) for 6 hrs, 1 mM H_2_O_2_ for 6 hrs, 1 mM AZC for 15 hrs. To determine sensitivity to HS, cells seeded at a density of 4 × 10^5^ were placed in a 43 °C incubator for 0-60 min and allowed to recover in a 37°C incubator for 48 hrs. To assess HS-induced protein aggregation, cells seeded at a density of 4 × 10^6^ were placed in a 43 °C incubator for 0-60 min and immediately harvested to isolate the soluble and insoluble protein fractions. To assess the heat sensitivity of yeast cells, mutant and WT strains were grown in YPD medium at 30 °C to mid-log phase (OD_600_ = 0.6-0.7). For spot tests, cells in liquid medium with an OD_600_ of 0.2 were spotted onto YPD agar plates, along with five spots of serial 5-fold dilutions. Plates were incubated at 37°C and 39°C for 3 days. Control plates were incubated at 30°C.

#### Pharmacological inhibition of biological processes

All experiments were performed with human WT and KO cells. To determine the effect of Hsp90 inhibition on the cell cycle and cell morphology, cells seeded at a density of 4 × 10^5^ were treated with GA (0-1000 nM) and PU-H71 (0-1000 nM) for 24 hrs. For cell death analyses, cells seeded at a density of 4 × 10^5^ were treated with GA (0-1000 nM) for 48 hrs and Novobiocin (0-1000 *μ*M) for 24 hrs. To assess the effect of GA on the degradation of Hsp90 clients or aggregation of intracellular proteins, cells seeded at a density of 2 × 10^6^ were treated for 24 hrs with either 0-400 nM or 0-750 nM GA accordingly. To assess the effect of Hsp70 inhibition on cell cycle and cell morphology, cells seeded at a density of 4 × 10^5^ were treated with JG-98 (0-2.5 *μ*M) for 24 hrs. To assess the effect of Hsp90 inhibition on proteasomal activity and the stability of 19S RP-related proteins, cells seeded at a density of 2 × 10^6^ were treated with GA (0-1000 nM) or JG-98 (0-5 *μ*M) or both for 24 hrs. Cells seeded at a density of 2-4 × 10^5^ were treated with proteasomal inhibitors MG132 and bortezomib for 24 hrs and 48 hrs respectively, to determine the effect of proteasome inhibition on cell death and cell morphology. To analyze the effect of the inhibition of ubiquitination on cell death and cell morphology, cells seeded at a density of 4 × 10^5^ were treated with PYR41 (0-30 *μ*M) for 48 hrs.

Proteotoxic challenge experiments were performed by treating cells seeded at a density of 4 × 10^5^ with 500 nM GA for 6 hrs, followed by a 24 hrs treatment with either 10 *μ*M MG132 or 500 nM Bortezomib. Consequent cell death was analyzed by flow cytometry. Alternatively, to analyze cell death, cells seeded at a density of 4 × 10^5^ were heat shocked at 43 °C for 30-45 min before a 24 hrs treatment with MG132 (0-20 *μ*M) at 37 °C. Cycloheximide (CHX) chase experiments were performed by treating cells seeded at a density of 2 × 10^6^ with 50 *μ*g/ml CHX for 0-8 hrs and harvested for immunoblot analyses. To visualize the accumulation of ubiquitinated proteins during proteasomal inhibition, cells seeded at a density of 1 × 10^6^ were treated with MG132 (0-10 *μ*M) for 24 hrs and harvested for immunoblot analyses.

Functional Hop rescue experiments were performed by transfecting KO cells seeded at a density of 5 × 10^5^ with HA-tagged WT Hop. Transfections utilized Lipofectamine 3000 reagent according to the manufacturer’s protocol. At 24 hrs post-transfection, cells were treated with MG132 (0-20 *μ*M) or GA (0-1000 nM) for 24 hrs before the rescue effect on cell cycle and cell death was analyzed.

#### Flow cytometry

##### Annexin V-PI staining

Human cells seeded at a density of 5 × 10^5^ were harvested by trypsinization, washed in phosphate buffered saline (PBS) and resuspended in 100 *μ*l annexin V-binding buffer (10 mM HEPES pH 7.4, 150 mm NaCl, 2.5 mm CaCl_2_). Annexin V-FITC (5 *μ*l) and propidium iodide (PI; 2.5 *μ*g/ml) were added to the cells and incubated for 15-20 min at 4 °C. Cells were analyzed by flow cytometry in a FACSCaliber (BD Biosciences) using CellQuest Pro software and data were analyzed by FlowJo. In the dot plot analyses, the first quadrant represented healthy, unstained cells; the second quadrant represented early apoptotic cells with only annexin V-FITC staining; the third quadrant represented late apoptotic cells with both annexin V-FITC and PI staining; and the fourth quadrant represented necrotic cells with PI staining.

##### Cell death assays

Human cells were resuspended in 100-200 *μ*l of PBS containing PI (2.5 *μ*g/ml) for 15-20 min at room temperature (RT) before flow cytometry analysis.

##### Cell cycle analyses

Cells were harvested as outlined above. Cells were fixed with 70% ice-cold ethanol, washed, treated with 100 *μ*g/ml RNase A at RT for 5 min, then incubated with 50 *μ*g/ml PI for 15-20 min at RT before flow cytometry analysis. Apoptotic cells were identified by the quantitation of the SubG0 cell population. S and G2/M phase cell cycle arrests were calculated by subtraction of S and G2/M phase cells of untreated control from the inhibitor-treated sets.

For all flow cytometry analyses, a minimum of 10,000 cells were analyzed for each sample.

#### Cell proliferation assay

Human cells seeded at a density of 4 × 10^4^ were grown for 24, 48 or 72 hrs after which, 100 *μ*g/ml MTT (3-(4,5-dimethylthiazol-2-yl)-2,5-diphenyltetrazolium bromide;) was added to the culture medium for 3 hrs. The culture medium was removed and the formazan crystals were dissolved in DMSO, the OD was measured at 550 nm by a plate reader (Tecan Sunrise).

#### Biochemical fractionation of soluble and insoluble proteins

To analyze the polyglutamine (polyQ) protein aggregation within cells, cells seeded at a density of 7 × 10^5^ were transfected with pEGFP-Q74 and pEGFP-Q23 plasmids by PEI. At 48 hrs post-transfection, cells were harvested for biochemical fractionation. Alternatively, at 24 hrs post-transfection, cells were treated with GA (0-1000 nM) for 24 hrs. For the comparative analysis between WT and KO cells, soluble-insoluble fractions were biochemically separated from 4 × 10^6^ cells for each genotype as described previously (Hjerpe et al., 2016). Briefly, cells were lysed in a lysis buffer (20 mM Tris-HCl pH 7.4, 2 mM EDTA, 150 mM NaCl, 1.2% sodium deoxycholate, 1.2% Triton-X100, 200 mM iodoacetamide, protease inhibitor cocktail [PIC]), sonicated (low power, 3 cycles of 10 s pulses), and centrifuged at 16,100 × *g* for 20 min. The supernatant was collected as the soluble fraction. The precipitate (insoluble fraction) was washed 5-6 times with PBS and solubilized in NuPAGE protein sample buffer (Life Technologies, cat no. NP0008) containing 10 mM DTT. Both biochemical fractions were analyzed by immunoblotting.

#### *In vitro* proteasomal activity assay

Human cells were harvested and washed. Cell pellets were resuspended in lysis buffer (25 mM Tris-HCl pH 7.4, 250 mM sucrose, 5 mM MgCl_2_, 1% NP-40, 1 mM DTT, 1 mM ATP) and incubated for 10-15 min on ice. Samples were centrifuged at 16,100 × *g* for 20 min and supernatants were collected for the proteasomal activity assay. Yeast cells were collected, washed with H_2_O and lysed mechanically with glass beads (3 × 30 s pulses) in lysis buffer (10 mM Tris-HCl pH 7.4, 50 mM NaCl, 10 mM MgCl_2_, 1 mM EDTA, 20% glycerol, 1 mM DTT, 1 mM ATP). Samples were centrifuged at 16,100 × *g* for 20 min and supernatants were collected for the proteasomal activity assay. Equal amounts of protein (25-50 *μ*g) for each sample was diluted in proteasomal reaction buffer (50 mM Tris-HCl pH 7.4, 5 mM MgCl_2_, 1 mM DTT, 1 mM ATP) in a 96-well opaque bottom white plate and 50 *μ*M Suc-LLVY-AMC was added to each well. AMC fluorescence was measured at 460 nm wavelength for the duration of 5-60 min, with an excitation at the wavelength of 380 nm. Purified proteasomes (1 *μ*g) were diluted in proteasomal reaction buffer and activity was measured with Suc-LLVY-AMC. Alternatively, the Proteasome Activity Assay Kit (Abcam) was used according to the manufacturer’s instructions to measure proteasomal activity. All fluorescence measurements were recorded using a plate reader (Cytation 3, BioTek).

#### *In vivo* proteasomal activity assay

Cells seeded at a density of 5 × 10^5^ were transfected with Ub-M-GFP (stable GFP) and Ub-R-GFP (degradation prone GFP) plasmids by PEI. At 48 hrs post-transfection, cells were harvested by trypsinization and GFP positive cells were measured by flow cytometry (Dantuma et al., 2000). The *in vivo* proteasomal activity was measured as the % Ub-M-GFP positive cells minus the % Ub-R-GFP positive cells.

#### Protein expression and purification

##### Hsp90β, Hsp90α and mutants

Human Hsp90s were expressed and purified as described previously (Nguyen et al., 2017; Moran Luengo et al., 2018). Hsp90s were expressed from the IPTG-inducible pCA528 bacterial expression vector (Andreasson et al., 2008) as fusion proteins with an N-terminal 6× His-SUMO-tag in the BL21 (DE3) Star, pCodonPlus or Rosetta (DE3) *E. coli* strains. Bacteria were harvested by centrifugation, resuspended in lysis buffer (40 mM HEPES-KOH pH 7.5, 100 mM KCl, 5 mM MgCl_2_, 10% glycerol, 4 mM β-mercaptoethanol (β-ME), 5 mM PMSF, 1 mM pepstatin A, 1 mM aprotinin, 1 mM leupeptin) and lysed by a microfluidizer EmulsiFlex-C5 or a French press. Clarified lysates were incubated with a Ni-IDA affinity matrix and protein was eluted with lysis buffer containing 250 mM imidazole. To remove the SUMO-tag, Ulp1-SUMO protease was added to the eluted protein and dialyzed against lysis buffer containing 20 mM KCl overnight at 4 °C. The dialyzed protein was loaded onto a Ni-IDA affinity matrix, followed by anion exchange chromatography (ResourceQ™; GE Healthcare Lifesciences) using a linear gradient of 0.02-1 M KCl. Eluted fractions of Hsp90 were further purified by size exclusion chromatography on a Superdex 200 (GE Healthcare Lifesciences) in storage buffer (40 mM HEPES-KOH pH 7.5, 50 mM KCl, 5 mM MgCl_2_, 10% glycerol, 4 mM β-ME).

##### Hsp70 and mutant

WT human Hsp70 and the substrate binding mutant (V435F) were expressed in BL21 (DE3) Rosetta cells and purified as an N-terminal 6× His-SUMO fusion protein (Moran Luengo et al., 2018). Bacterial cells were resuspended in lysis buffer (20 mM Tris-HCl pH 7.9, 100 mM KCl, 1 mM PMSF) and lysed by microfluidizer EmusiFlex C5 or a French press. Clarified lysates were loaded onto Ni-IDA affinity matrixes. The affinity matrix was washed first with lysis buffer (without PMSF), then with high salt buffer (20 mM Tris-HCl pH 7.9, 1 M KCl), and again with lysis buffer. The affinity matrix was washed with ATP-buffer (40 mM Tris-HCl pH 7.9, 100 mM KCl, 5 mM MgCl_2_, 5 mM ATP) to elute bound proteins. Hsp70 was eluted with 1 column volume of elution buffer (40 mM Tris-HCl pH 7.9, 100 mM KCl, 250 mM imidazole). Eluted proteins were dialyzed O/N against dialysis buffer (40 mM HEPES-KOH pH 7.6, 10 mM KCl, 5 mM MgCl_2_) in the presence of Ulp1. The dialyzed sample was loaded onto a Ni-IDA affinity matrix and the flow-through containing the Hsp70 protein was subjected to a Resource™ Q anion exchange column. Hsp70 protein was eluted in elution buffer (40 mM HEPES-KOH pH 7.6, 1 M KCl, 5 mM MgCl_2_, 10 mM β-ME, 5% glycerol) and further dialyzed against storage buffer (40 mM HEPES-KOH pH 7.6, 50 mM KCl, 5 mM MgCl_2_, 10 mM β-ME, 10% glycerol).

##### Hop and Apg2

Hop and Apg2 proteins were expressed in BL21 (DE3) Rosetta cells (Merck KGaA, Darmstadt, Germany) and purified as an N-terminal 6× His-SUMO fusion protein as described previously (Moran Luengo et al., 2018). Bacterial cells were resuspended in lysis buffer (40 mM Tris-HCl pH 7.9, 100 mM KCl, 5 mM ATP, 8 mg/l Pepstatin, 10 mg/l Aprotinin, 5 mg/l Leupeptin), and lysed by a microfluidizer EmulsiFlex-C5. The clarified lysates were loaded onto Ni-IDA affinity matrixes. Protein was eluted with buffer (40 mM Tris-HCl pH 7.9, 100 mM KCl) containing 300 mM imidazole. Subsequently, buffer exchange was performed using a HiPrep 26/10 Desalting Column (GE Healthcare Lifesciences). Ulp1-SUMO protease was added to the eluted protein and the solution was incubated O/N in the presence of 5 mM ATP. The protein was purified using a HiLoad 16/600 Superdex 200 gel filtration column (GE Healthcare Lifesciences). Gel filtration eluents were further purified using anion-exchange chromatography on a Resource™ Q column (GE Healthcare Lifesciences) for Apg2, and POROS 20HQ column (Thermo Fisher Scientific) for Hop. Both proteins were eluted with a linear KCl gradient (0.01−1 M).

##### DnaJB1

Human DnaJB1 was expressed and purified as a 6× His-SUMO fusion protein as described previously (Malakhov et al., 2004).

##### Firefly luciferase

*Photinus pyralis* firefly luciferase was expressed and purified as described previously (Rampelt et al., 2012). Firefly luciferase expressing XL10 Gold bacterial cells (Stratagene, US) were resuspended in lysis buffer (50 mM Na_x_H_y_PO_4_ pH 8.0, 300 mM NaCl, 10 mM β-ME, protease inhibitors, 10 *μ*g/ml DNase) and lysed by microfluidizer EmulsiFlex-C5. Clarified lysate was applied to Ni-IDA affinity matrix beads and firefly luciferase was eluted with elution buffer (50 mM Na_x_H_y_PO_4_ pH 8.0, 300 mM NaCl, 250 mM Imidazole, 5 mM β-ME). Purified firefly luciferase was dialyzed O/N against dialysis buffer (50 mM Na_x_H_y_PO_4_ pH 8.0, 300 mM NaCl and 10 mM β-ME, 10% glycerol).

##### GST-UBL

GST-UBL was expressed and purified as described previously (Besche and Goldberg, 2012). GST-UBL was induced using 0.1% (w/v) L-arabinose in the Rosetta (DE3) strain. Rosetta (DE3) cells were resuspended in GSH binding buffer (1× PBS, 10 mM MgCl_2_, 1 mM DTT, PIC) and lysed by French press. The clarified lysate was incubated with GSH-agarose affinity matrix and GST-UBL was eluted with an elution buffer (100 mM Tris-HCl pH 8, 100 mM NaCl, 1 mM DTT, 20 mM reduced GSH). Eluted protein was dialyzed twice against storage buffer (25 mM HEPES-KOH pH 7.4, 40 mM KCl, 5 mM MgCl_2_, 10% glycerol, 1 mM DTT).

##### 10× His-tagged-UIM

Expression of His-tagged-UIM was induced by IPTG in the Rosetta (DE3) bacterial strain. Bacterial cells were resuspended in lysis buffer (25 mM HEPES-KOH pH 7.4, 500 mM NaCl, 1 mM DTT, 0.025% NP-40, 20 mM imidazole) and lysed by French press. The clarified lysate was incubated with Ni-NTA affinity matrix and His-tagged-UIM was eluted with an elution buffer (25 mM HEPES-KOH pH 7.4, 500 mM NaCl, 1 mM DTT, 0.025% NP-40, 500 mM imidazole). Eluted protein was dialyzed twice against the storage buffer (Besche and Goldberg, 2012).

#### 26S/30S proteasome purification from human cells

Proteasomes were purified from mammalian cells as described previously (Besche and Goldberg, 2012). Human cells were cultured in 15 cm cell culture dishes until 85-90% confluency was reached. Cells were harvested, resuspended in lysis buffer (25 mM HEPES-KOH pH 7.4, 40 mM KCl, 5 mM MgCl_2_, 10% glycerol, 1 mM DTT, 1 mM ATP) and lysed by sonication (3 cycles of 30 s pulses). The lysates were clarified by centrifugation and the supernatants were filtered through 0.45 μm membranes. Clarified lysates were incubated with 1 mg GST-UBL (final concentration of 0.1-0.2 mg/ml) and GSH-agarose beads. The mixture was loaded onto a column and washed with 40 column volumes of lysis buffer. The proteasome was eluted in two rounds of 250 *μ*l of 10× His-tagged UIM (≥2 mg/ml) with a 15 min incubation before elution at 4 °C. The eluted proteasomal fraction was further applied on Ni-IDA affinity matrix to remove excess 10× His-tagged UIM. Purified proteasomes were stored at −80 °C.

#### *In vitro* luciferase refolding assay

Firefly luciferase was diluted to 100 *μ*M in refolding buffer (25 mM HEPES-KOH pH 7.6, 100 mM KOAc, 10 mM Mg(OAc)_2_, 2 mM ATP, 5 mM DTT) containing 1 *μ*M Hsp70, DnaJB1 and Apg2 (Hsp70:DnaJB1:Apg2; 2:1:0.5) and 1 *μ*M Hsp90α, when indicated. Recombinant human Hop was added in the indicated experimental sets in different concentrations (Hsp90α:Hop = 1:0.25, 1:0.5, 1:1).The firefly luciferase was heat denatured at 42 °C for 10 min and refolding was allowed at 37 °C for 10-45 min. Luciferase activity of the reaction mixture was measured in assay buffer (100 mM K-phosphate buffer pH 7.6, 25 mM glycylglycine, 100 mM KOAc, 15 mM Mg(OAc)_2_, 5 mM ATP) by adding the substrate, luciferin (final concentration 100 *μ*M). Measurements were obtained using a luminometer plate reader (Berthold Technologies, XS3 LB930; Mikrowin Software 2010).

#### *In vivo* luciferase refolding assay

##### Human cells

Cells (2 × 10^4^) transfected with the luciferase expression vector pC7L were resuspended in 100 *μ*l of cell culture medium. Cells underwent HS at 42-43 °C for 15 min to denature luciferase, followed by incubation at 37 °C for 1-3 hrs for the refolding of luciferase. Cells were harvested by centrifugation and lysed with Passive Lysis Buffer (Promega). Cell extract (10 *μ*l) was mixed with an equal volume of firefly luciferase assay substrate from the Dual Luciferase detection kit (Promega) and the luciferase luminescence signals were measured by the Chameleon bioluminescence plate reader (Noki tech.). To determine the impacts of Hsp70 and Hsp90 inhibitors on luciferase refolding, 2 *μ*M of JG-98 or 1 *μ*M of GA was added to pC7L transfected cells 1 hr prior to HS. HS and luciferase refolding were performed in complete cell culture medium containing the same concentrations of JG-98 and GA. The luciferase activity of cells not subjected to HS was set to 100%. Alternatively, cells were co-transfected with pC7L and either WT or TPR mutant Hop. At 48 hrs post-transfection a luciferase refolding assay was performed as described above. However, luciferase was heat denatured at 44 °C for 20 min.

##### Yeast cells

pLG/Luc transformants were grown O/N in galactose-containing minimal medium at 30 °C. Cells were diluted and grown to mid-log phase (OD_600_ = 0.6-0.7), washed with water and resuspended in minimal medium containing glucose and 100 *μ*g/ml CHX; OD_600_ was adjusted to 0.4 for each sample. 100 *μ*l samples of yeast cells were subjected to HS at 44 °C for 10-30 min to heat-denature luciferase, the refolding of luciferase was allowed by incubation at 30 °C for 1-3 hrs. Whole cell luciferase activity was measured by mixing 50 *μ*l of processed yeast cells with equal volumes of 100 mM Na-citrate buffer (pH 5.0) and 1 mM D-luciferin, before luminescence was measured with a bioluminescence plate reader. The luciferase activity of cells not subjected to HS was set to 100%.

#### Immunoprecipitation (IP)

##### Hsp90α, Hsp90β, HA-Hsp70, and FLAG-Hsp90α IP

Human cells were resuspended in receptor buffer (10 mM Tris-HCl pH 7.5, 50 mM NaCl, 1 mM EDTA, 1 mM DTT, 10% glycerol, 10 mM Na-molybdate, PIC) and lysed by sonication (30-40 cycles of 30 s) using a Bioruptor^®^ sonicator (Diagenode). For Hsp90α or Hsp90β IP, 1 mg of clarified cell extract was mixed with 5 *μ*g anti-Hsp90α (9D2) or Hsp90β (H90-10) antibodies. For HA-Hsp70 or FLAG-Hsp90α-IP, ~2 mg of cell extract was mixed with either 5-8 *μ*g anti-HA (HA.11) or anti-FLAG (M2) antibodies.

##### 20S CP IP

Human cells were lysed in a lysis buffer (25 mM Tris-HCl pH 7.4, 250 mM Sucrose, 5 mM MgCl_2_, 1% NP-40, 1 mM DTT, 1 mM ATP, PIC), 2 mg of clarified cell extract was mixed with 5 *μ*g anti-20S CP-specific antibodies. For all experiments, IPs were incubated O/N at 4 °C on a rotating wheel before 50 *μ*l of Dynabeads™-ProteinG (Thermo Fisher Scientific) was added and incubated for 3 hrs at 4 °C. The Dynabeads were washed and boiled with the NuPAGE protein sample buffer containing 10 mM DTT. The eluates were collected, separated by SDS-PAGE (7.5%-10%) and visualized using immunoblotting.

#### Native-PAGE analysis

To determine the constituents of proteasomal complexes in total cell lysates, human cells (lysis buffer: 25 mM Tris-HCl pH 7.4, 250 mM sucrose, 5 mM MgCl_2_, 1% NP-40, 1 mM DTT, 1 mM ATP, PIC) and yeast cells (lysis buffer: 10 mM Tris-HCl pH 7.4, 50 mM NaCl, 10 mM MgCl_2_, 1 mM EDTA, 20% glycerol, 1 mM DTT, 1 mM ATP, PIC) were lysed. Equal amounts of total cell extracts (50-75 *μ*g) or purified proteasomes (2 *μ*g) were separated by 4% Tris-Glycine Native-PAGE. For all the samples, Native-PAGE was performed for 3.5-4 hrs at 100-110 mV in a 4 °C cold room. Separated proteins were transferred onto nitrocellulose membrane and visualized with proteasomal subunit-specific primary antibodies.

#### *In vitro* protein-protein interaction assay

5 *μ*g of purified Hsp90α or Hsp90β protein was mixed with 7.5 *μ*g of purified Hsp70 (V435F) in association buffer (10 mM Tris-HCl pH 7.5, 50 mM NaCl, 1 mM EDTA, 1 mM DTT, 0.05% Triton-X100, 10% glycerol, 10 mM Na-molybdate, PIC) in the presence or absence of 1 mM ATP. The mixtures were incubated for 2 hrs at 30 °C and the association of Hsp90 and Hsp70 was determined by a co-IP. Briefly, 5 *μ*g anti-Hsp90α (9D2) or Hsp90β (H90-10) antibodies along with 25 *μ*l of Dynabeads™-ProteinG were added to the protein mixtures and incubated O/N at 4°C. Equal amounts of normal IgG were used for the corresponding control of co-IP. Dynabeads were washed and boiled with the NuPAGE protein sample buffer containing 10 mM DTT, and elutes were collected. Further SDS-PAGE and immunoblotting were performed with eluted protein samples, using purified proteins as inputs.

#### Mass spectrometry

##### Hsp90β IP-MS sample preparation

Three biological replicates of WT and KO cells from HEK293T and HCT116 backgrounds were lysed in lysis buffer (20 mM HEPES-KOH pH 7.5, 50 mM KCl, 5 mM MgCl_2_, 20 mM Na-molybdate, 1% NP40, protease inhibitors, phosphatase inhibitors). Clarified protein lysates (500 μg) were diluted in a total volume of 140 *μ*l of incubation buffer (20 mM HEPES-KOH pH 7.5, 50 mM KCl, 5 mM MgCl_2_, 20 mM Na-molybdate, 0.01% NP40). Hsp90-IPs were generated by incubating cell lysates with 15 *μ*g of anti-Hsp90β and 20 μg anti-Hsp90α antibodies. Control IPs were performed with equal amounts of normal mouse IgG and normal rat IgG2a for 4 hrs at 4 °C on a rotating wheel. Protein A-agarose (40 *μ*l of a 50% slurry) was added to the IP mixtures and incubated for an additional 2 hrs. Beads were washed and proteins eluted with 60 *μ*l FASP lysis buffer (100 mM Tris-HCl pH 7.5, 4% SDS, 10 mM TCEP) at 95 °C for 5 min. The samples were digested by the FASP method as described previously (Wisniewski et al., 2009). Resulting peptide mixtures were desalted on Waters SEP-PAK C18 micro elution plates and eluted with 100 *μ*l of 40% acetonitrile, 0.1% formic acid. Eluates were separated in 6 fractions on SOLA SCX SPE columns (Thermo Fisher) using an increasing concentration of ammonium acetate (50, 75, 125, 250, 500, 1000 mM) in 20% acetonitrile and 0.5% formic acid. The dried peptides were resuspended in 25 *μ*l solvent A (2% acetonitrile, 0.1% formic acid) and 5 *μ*l samples were used for LC-MS/MS analysis.

##### Hop (HA)-IP-MS sample preparation

Two biological replicates of HEK293T Hop KO cells were transfected with WT or TPR double mutant (K8, 229A) Hop mammalian expression plasmids. Empty vector transfected cells were considered a control for this experiment. At 48 hrs post-transfection, cells were lysed by sonication in receptor buffer. Clarified cell lysates (2 mg) were incubated with 10 μg anti-HA antibodies O/N at 4 °C to IP exogenous HA-tagged WT or TPR double mutant Hop. Dynabeads™-Protein G was added to the incubations and proteins were eluted as described above. Eluted proteins were separated by 12% 1D-SDS-PAGE over a distance of 4.0 cm. After Coomassie blue staining, entire lanes were excised and underwent in-gel reduction and alkylation with chloroacetamide, and then digested with trypsin as described previously (Wilm et al., 1996). Extracted peptide mixtures were subjected to LC-MS/MS analysis.

##### Whole cell proteome MS sample preparation

Three biological replicates of WT and KO HEK293T and HCT116 cells were lysed and the proteins digested according to a modified version of the iST protocol (Kulak et al., 2014). Cells (5 × 10^6^) were lysed in 250 *μ*l modified iST buffer (100 mM Tris-HCl pH 8.6, 1% sodium deoxycholate, 10 mM DTT, protease inhibitors, phosphatase inhibitors) and heated at 95 °C for 5 min. The lysates were diluted 1:1 with 4 mM MgCl_2_, benzonase nuclease was added and incubated for 15 min at RT. EDTA (3 mM) and 30 mM chloroacetamide were added for 45 min at 25 °C in the dark to alkylate cysteine residues. Samples were digested first with 2.5 μg of trypsin/Lys-C mix for 1 hr at 37 °C, followed by a second enzyme addition (1.25 μg trypsin/LysC) for 1 hr at 37 °C. To extract deoxycholate, 2 volumes of ethyl acetate and 1% TFA were added to 1 volume of lysate, the mixture was vortexed for 2 min and centrifuged. 100 *μ*l of aqueous fractions were loaded onto equilibrated SOLA SCX SPE columns (Thermo Fisher) prefilled with 450 *μ*l SCX0 buffer (20% acetonitrile, 0.5% formic acid) and centrifuged. The columns were washed once with 300 *μ*l of ethyl acetate and 0.5% TFA solution, and twice with 300 *μ*l solvent A. The peptide mixtures were sequentially eluted with 200 *μ*l SCX125 buffer (20% acetonitrile, 0.5% formic acid, 125 mM ammonium acetate), 200 *μ*l SCX500 buffer (20% acetonitrile, 0.5% formic acid, 500 mM ammonium acetate), and finally with 200 *μ*l basic elution buffer (80% acetonitrile, 19% H_2_O, 0.25% NH_3_). The dried fractions were resuspended in 100 *μ*l solvent A and 5 *μ*l solutions were used for LC-MS/MS analysis.

##### Purified proteasome MS sample preparation

Each sample preparation used 15-25 μg purified proteasomes. Chloroacetamide was added at the final concentration of 5 mM. The samples were diluted 1:1 with 8 M urea and incubated at RT for 30 min to alkylate cysteine residues. Endoproteinase LysC (0.2 *μ*g) was added to the mixtures and the digestion was carried out for 2 hrs at 37 °C. The resulting solutions were diluted with 50 mM ammonium bicarbonate buffer and 0.5 μg of sequencing grade trypsin (Promega) was added before the samples were incubated at 37 °C O/N. Digested samples were acidified with formic acid, and half of each digested sample was desalted on Waters C18 microelution plates by centrifugation. Peptides were eluted with 100 *μ*l of 40% acetonitrile and 0.1% formic acid solution. Dried samples were resuspended in 25 *μ*l 0.05% TFA for LC-MS/MS analysis.

##### General LC-MS/MS analysis

Samples were analyzed on a Fusion orbitrap trihybrid mass spectrometer, interfaced via a nanospray source to a Dionex RSLC 3000 nano HPLC system (Thermo Fisher Scientific, Bremen, Germany). Extracted peptide mixtures were separated on a custom packed reversed-phase C18 nanocolumn (75 μm ID × 40 cm, 1.8 μm particles, Reprosil Pur, Dr. Maisch) with a gradient from 5 to 55% acetonitrile in 0.1% formic acid for 120 min (for IP samples: same gradient in 35 min). Full MS survey scans were performed at 120,000 resolution. All survey scans were internally calibrated using the 445.1200 background ion mass. Using a data-dependent acquisition controlled by Xcalibur 2.1 software (Thermo Fisher), a maximum number of multi-charged precursor ions was selected for tandem MS analysis within a maximum cycle time of 3 s. Selected precursor ions were fragmented by Collision-Induced Dissociation (CID) and analyzed in the linear ion trap with an isolation window of 1.6 m/z. Selected ions were then dynamically excluded from further selection during 60 s.

#### Phase contrast and fluorescence microscopy

Cellular morphology was analyzed using an inverted light microscope (Olympus CK2) at 10× magnification and phase contrast images were captured with a Dino-lite camera using DinoXcope software. Cells were seeded on glass coverslips and transfected with Q74-EGFP and Q23-EGFP plasmids. At 48 hrs post-transfection, cells were fixed with 4% paraformaldehyde and mounted on glass slides using Mowiol mounting medium. EGFP positive cells were visualized and images were captured with a fluorescence microscope (Zeiss, Germany) at 20× magnification.

#### Negative staining transmission electron microscopy

Carbon-coated copper grids (200 or 300 mesh) were glow discharged for 20 s. Purified proteasomes were diluted in buffer (25 mM HEPES-KOH pH 7.4, 40 mM KCl, 10% [v/v] glycerol, 5 mM MgCl_2_, 1 mM ATP, 1 mM DTT) and 5 *μ*l of samples were loaded onto the grid surface and incubated for 30 s. Excess samples were blotted off using filter paper while holding the grid vertically. Two drops of (100 *μ*l each) uranyl acetate (2% in water) were prepared for each grid. Grids were incubated for 2 s on a uranyl acetate drop and on a second drop for 30 s to stain the grids. Processed grids were analyzed with a transmission electron microscope at 120 kV (Tecnai G2, FEI, Eindhoven, Netherlands).

#### Transcriptional activity assays

Human cells were seeded at a density of 6 × 10^4^ in phenol red-free DMEM medium containing 5% charcoal-treated FBS in 24-well cell culture plates. The cells were co-transfected with ERα, GR or PR mammalian expression plasmids (only for HEK293T cells), as well as both a luciferase reporter plasmid and a constitutive Renilla luciferase expression plasmid (pRL-CMV). At 48 hrs post-transfection, cells were lysed with Passive Lysis Buffer, and firefly and Renilla luciferase activities were measured using the Dual-Luciferase detection kit (Promega) with a bioluminescence plate reader. Renilla luciferase activity was used as transfection control.

#### Bioinformatic analyses

##### *In silico* protein homology modeling

Beginning with the apo conformation of HtpG (PDB ID: 2IOQ) (Shiau et al., 2006), homology models of the human Hsp90α and Hsp90β proteins were constructed using the I-TASSER structure-prediction web server (Roy et al., 2010). Briefly, monomeric structures were predicted according to the C-score and TM-score of Hsp90α (C-score = −1.53, TM-score = 0.53±0.15) and Hsp90β (C-score = −0.99, TM-score = 0.59±0.14). To generate dimer structures of Hsp90α and Hsp90β, full-length predicted monomeric structures of each protein were manipulated using the ClusPro protein-protein docking web server programmed in dimer mode (Kozakov et al., 2017). Hydrophobic interaction driven dimer structures, with low free energy, were selected. The accessibility of surface residues of the dimeric Hsp90α and Hsp90β were predicted by Swiss-PdbViewer 4.1.1 software.

##### Gene Ontology Enrichment analysis

Gene ontology enrichment analyses were performed with the Enricher web server. Significant biological processes were identified by lower P values, a higher number of common genes/proteins responsible for a biological process, and a high combined score provided by the Enricher web server.

##### Protein interaction network analysis

Fold change values (WT vs KO) of proteins from the whole cell MS analyses were analyzed as described previously (Echeverria et al., 2019). All nodes correspond to identified proteins. Cytoscape analyses were represented as heat maps of a log2 fold change of protein expressions. Hsp90-specific clients, chaperones and co-chaperones were classified using our own web server Hsp90Int.DB.

### QUANTIFICATION AND STATISTICAL ANALYSIS

#### General data analyses

Data processing and analyses were performed using GraphPad Prism7. Unless stated otherwise, the data shown are averages of at least three biological replicates with multiple corresponding technical replicates. Data are represented as mean values and the error bars represent the standard deviation of the mean. The statistical significance between the groups was analyzed by two tail Student's t-test with p < 0.05.

#### Mass spectrometric data analyses

Tandem MS data were processed by the MaxQuant software (ver.1.6.0.13) (Cox and Mann, 2008) incorporating the Andromeda search engine (Cox et al., 2011). The UniProt human reference proteome database (of October 2017) was used (71,803 sequences), supplemented with sequences of common contaminants. All identifications were filtered at 1% FDR at both the peptide and protein levels with default MaxQuant parameters. For the whole cell proteome and Hsp90-IP-MS experiments, from the relevant MS datasets only protein groups were kept for the subsequent analyses which met the criteria of at least 2 unique peptides, 4 peptides and 6 MS/MS counts. Protein quantitation was performed either with the iBAQ values (for the IP-MS analyses) or with the LFQ values (for the whole cell proteome and proteasomal analyses) (Cox et al., 2014). In whole cell proteome and Hsp90-IP-MS analyses, the log2 fold change > 0.4 or < −0.4 of a protein was considered as the biologically significant difference. Proteins that were detected in at least one of three biological replicates (for the whole cell proteome) were used in all further analyses.

In Hsp90-IP-MS analyses, the criteria for the validation of interactors was an average fold change of 8 (3 in log2 scale) or more, identified by MS/MS in all positive IP samples compared to control IP, and an adjusted p value < 0.05. The iBAQ values of interactors were normalized on the sum of the iBAQ of Hsp90βand Hsp90α.

In Hop (HA)-IP-MS analyses, the interactor validation criteria was an average fold change of 40 or more identified by MS/MS in WT Hop IP samples compared to the control IP and at least 3 unique peptides. Since chaperone, co-chaperone and proteasomal proteins were in our point of interest, we included three proteasomal components (PSMB1, PSMD7, and PSMD14) and three chaperone proteins (HSP90AB4P, HSP6, and HSP7) in our further analyses despite 2 unique peptides. The iBAQ values of interactors were normalized either on the abundance of WT or TPR mutant Hop depending on the experimental set. PCA and (O)PLS-DA analyses of the results from whole cell proteome and Hsp90-IP-MS experiments were done using the ropls package for R (Thevenot et al., 2015). The variable importance in projection (VIP) values were derived from the “Orthogonal Projections to Latent Structures Discriminant Analysis” (OPLS-DA) model and used to select variables of interest with a VIP cut-off of 1 (Galindo-Prieto et al., 2014).

#### Proteasomal activity analyses

Without having SDS in the reaction buffer, activity readouts using the reporter, suc-LLVY-AMC, could be considered the 26S/30S proteasomal activity. For steady-state proteasomal activity measurements, AMC fluorescence of WT samples for each time point was considered as 100% proteasomal activity and the corresponding proteasomal activity of KO samples was calculated. For the proteasomal activity rate measurements, AMC fluorescence of each time point was normalized with AMC fluorescence at 5 min; data were represented as the fold change while slopes were considered as the rate of proteasomal activity over time.

#### TEM image analyses

2D class averaging of different proteasomal structural projections were done by Scipion software. TEM micrographs were uploaded on Scipion software and analyzed using an optimized program. Lengths and diameters of different structures of the proteasome were determined by ImageJ software.

### DATA AND SOFTWARE AVAILABILITY

The mass spectrometry proteomics data are available through the ProteomeXchange Consortium (via the PRIDE partner repository) with the dataset identifier PXD012774 (not publicly accessible yet) and for a subset in Table S1.

## SUPPLEMENTAL INFORMATION

Supplemental Information includes seven figures and one table.

**Table S1. Quantitative MS Analyses of Hop (STIP1) Peptides in WT and KO1 HEK293T Cells, Related to Figure 1.**

### SUPPLEMENTAL FIGURE LEGENDS

**Figure S1.**
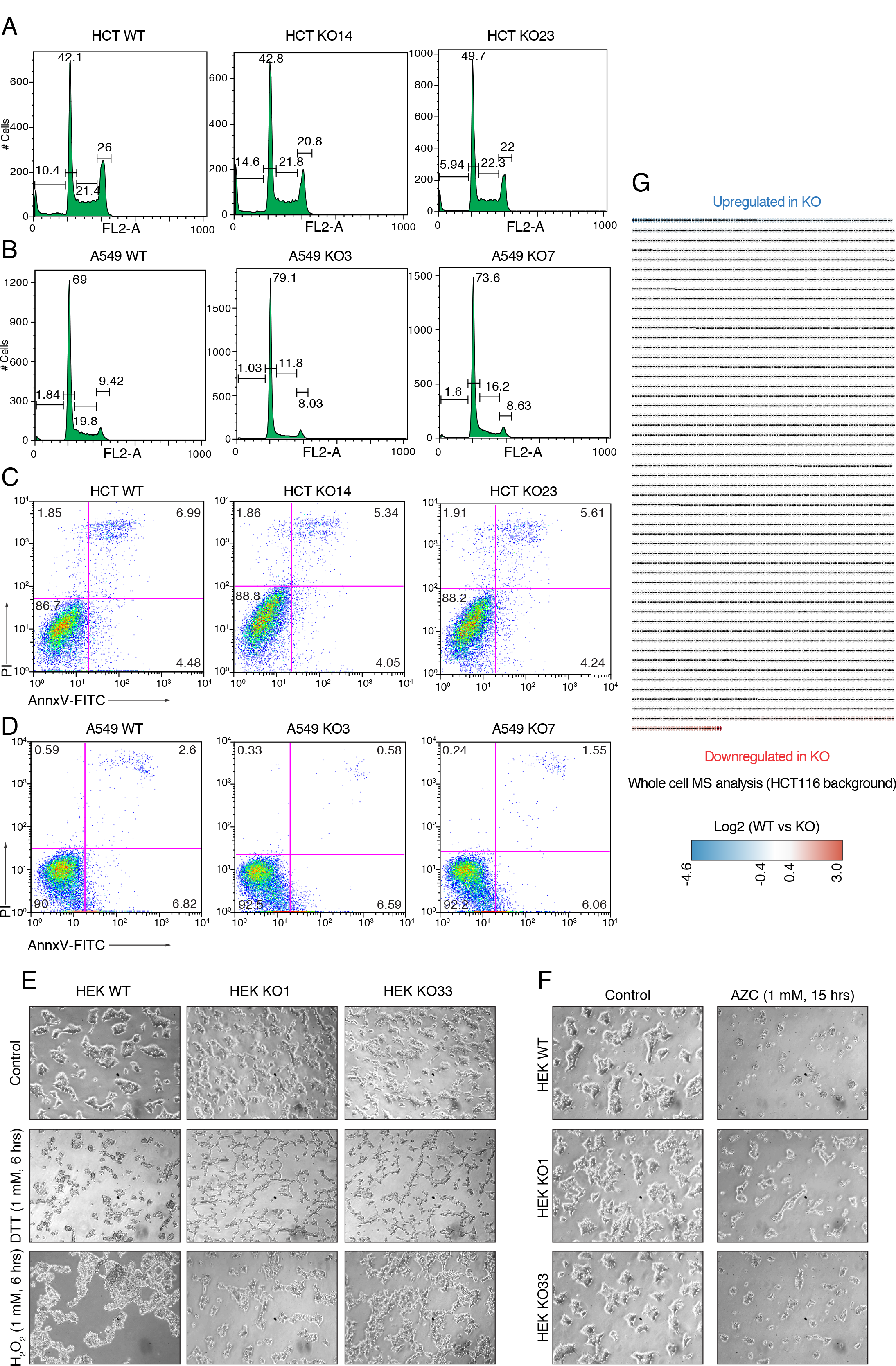
KO Cells Maintain Proteostasis and Are Not Hypersensitive to Proteotoxic Stresses, Related to Figure 1. (A, B) Flow cytometry histograms represent the cell cycle distributions of WT and KO cells. (C, D) Dot plots represent live/dead cells distributions visualized by annexin V-FITC (AnnxV-FITC) and PI staining using flow cytometry. (E, F) Phase contrast micrographs of WT and KO cells treated with DTT and H_2_O_2_ (panel E), and AZC (panel F). Vehicle-treated cells serve as a control. HEK, HEK293T. (G) Heat map of whole cell proteome changes in HCT116 cells (WT vs KO14).

**Figure S2.**
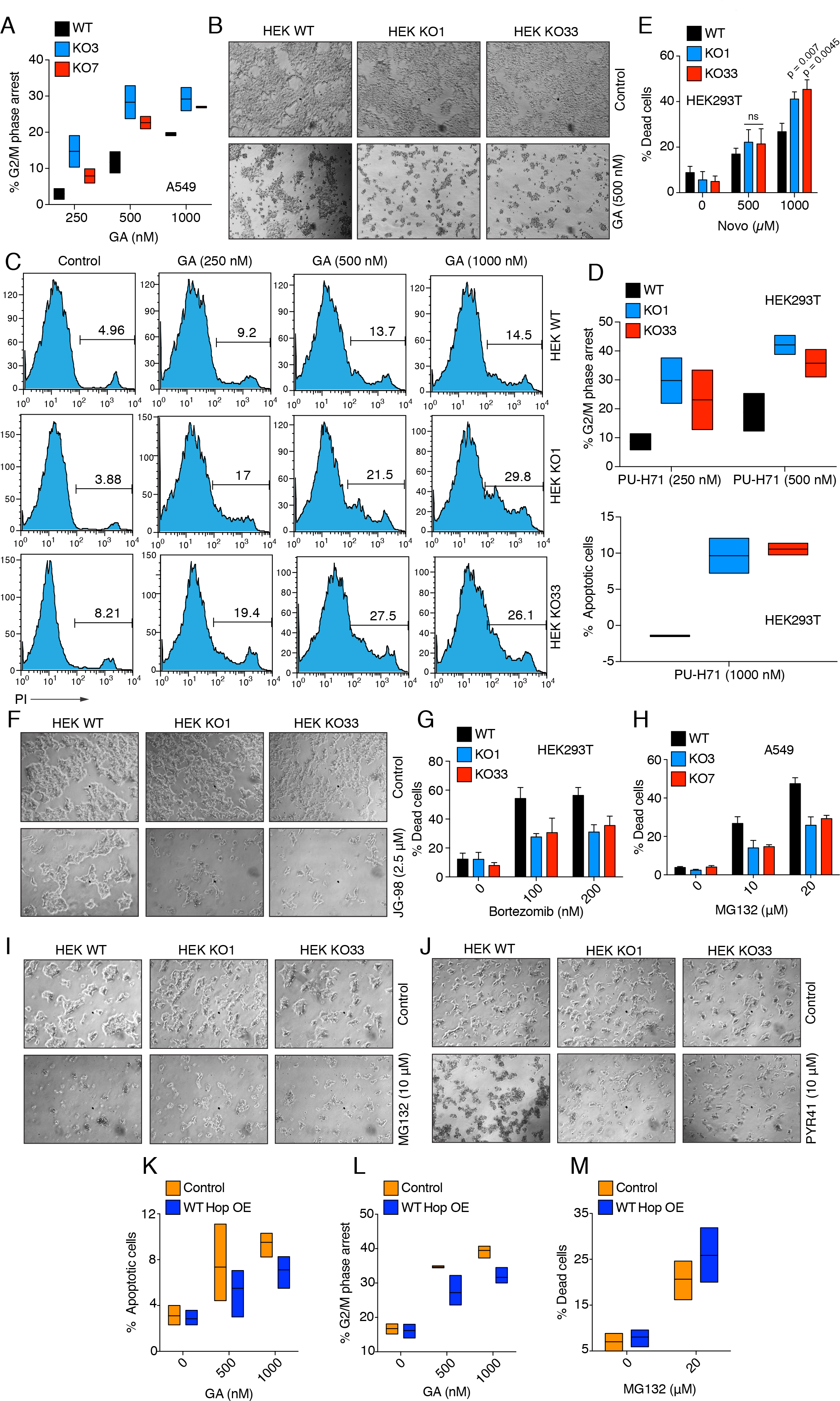
Inhibition of Hsp70 and Hsp90 or the Proteasome Have Differential Impacts on KO Cells, Related to Figure 2. (A) Flow cytometric analysis of the GA-induced G2/M phase cell cycle arrest with A549 cells treated with GA for 24 hrs (n=2). Data are represented as a box plot. (B) Phase contrast micrographs of WT and KO cells treated with GA for 24 hrs. Vehicle-treated cells serve as a control. (C) Flow cytometry histograms representing the impact of a treatment with GA for 48 hrs. The number over the linear gate in each plot indicates the % PI positive dead cells in the total analyzed cell population. (D) Flow cytometric analysis of the PU-H71-induced G2/M phase cell cycle arrest (top) and apoptosis (bottom) after 24 hrs of treatment (n=2). Data are represented as a box plot. (E) Flow cytometric analysis of cell death induced by novobiocin (Novo) of after 24 hrs of treatment. (F) Phase contrast micrographs of WT and KO cells treated with JG-98 for 24 hrs. Vehicle-treated cells serve as a control. (G, H) Flow cytometric analysis of bortezomib- and MG132-induced cell death after 48 and 24 hrs of treatment respectively. (I, J) Phase contrast micrographs of WT and KO cells treated with MG132 for 24 hrs and PYR41 for 48 hrs. Vehicle-treated cells serve as a control. (K, L) Flow cytometric analysis of GA-induced apoptosis and G2/M phase cell cycle arrest of KO HEK293T cells overexpressing (OE) full-length WT Hop after 24 hrs of treatment. Cells transfected with empty vector serve as a control. Data are represented as a box plot. (M) Flow cytometric analysis of MG132-induced cell death of KO HEK293T cells overexpressing (OE) full-length WT Hop. Data are represented as a box plot.

**Figure S3.**
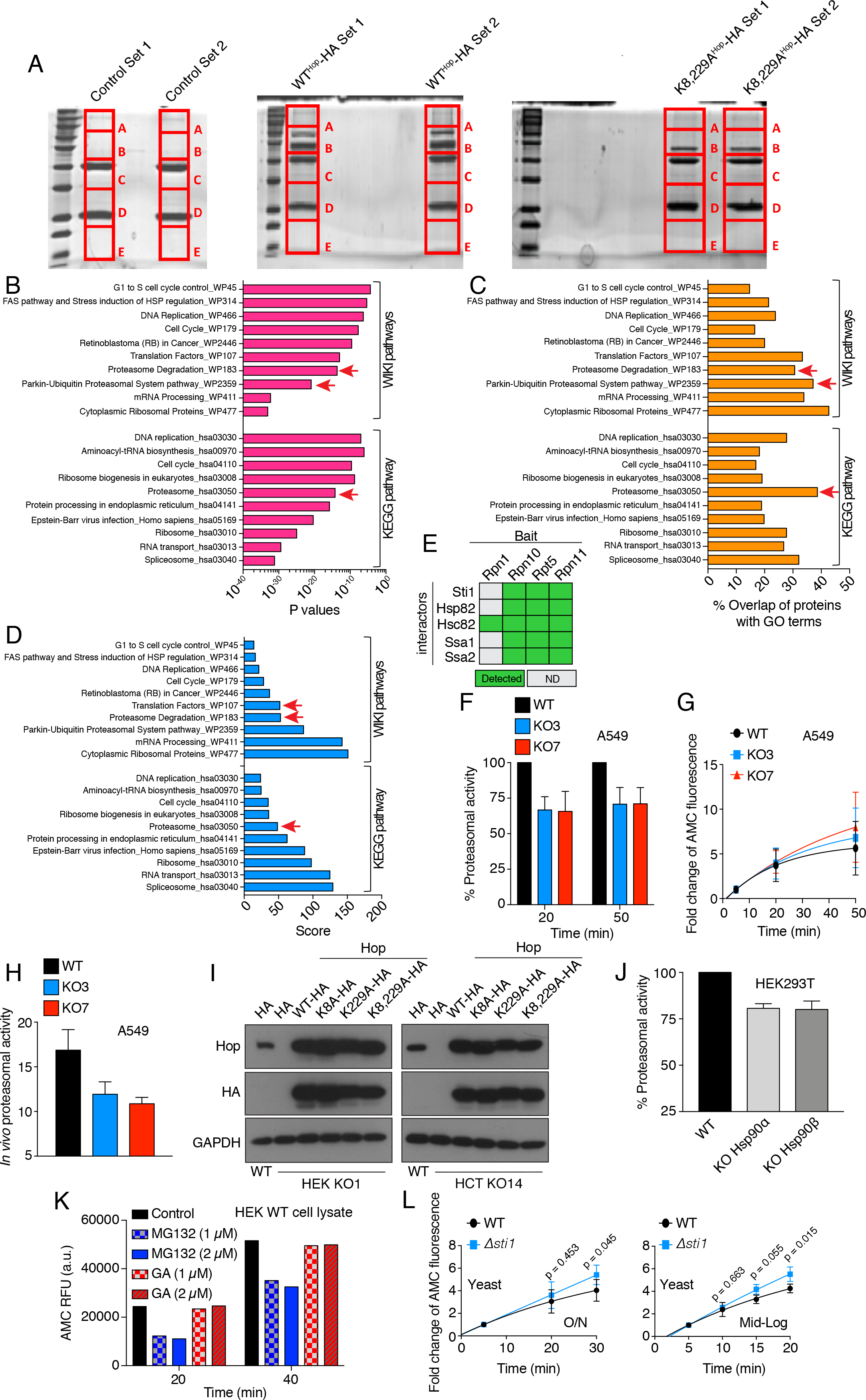
The Absence of the Hsp70-Hop-Hsp90 Ternary Complex Compromises Proteasomal Function in KO Cells, Related to Figure 3. (A) Anti-HA immunoprecipitations of WT and TPR double mutant (K8,229A) HA-tagged Hop overexpressed in HEK293T KO1 cells; cells transfected with empty vector were used as negative control. Two biological replicates for each sample were visualized by Coomassie staining following SDS-PAGE. Each lane was processed for LC-MS/MS analysis to identify Hop interactors. (B-D) GO term enrichment analyses of identified Hop interactors (see also Figure 3B). GO term enrichment analyses were done with the Enricher web server, and the top 10 KEGG and WIKI pathway-annotated biological processes are plotted according to the P values (panel B), % overlap of MS-identified proteins with annotated proteins having a known GO function (panel C), and a combined score provided by the Enricher web server (panel D). Proteasomal and ubiquitin-related pathways are indicated with red arrows. (E) Interaction matrix of components of the ternary molecular chaperone complex identified as interactors of proteasome components purified from yeast. Data were obtained from a previously published MS dataset of the affinity-purified proteasome (Guerrero et al., 2008). (F) *In vitro* steady-state proteasomal activity of extracts of WT and KO A549 cells. (G) Rate of proteasomal activity determined with the activity reporter suc-LLVY-AMC. (H) Flow cytometric determination of the *in vivo* proteasomal activity using the Ub-M-GFP and Ub-R-GFP reporter plasmids. (I) Immunoblots of overexpressed HA-tagged WT and TPR mutants of Hop. A representative immunoblot analysis for the quality control of cell extracts corresponding to those assayed for proteasomal activity in Figure 3H. (J) *In vitro* steady-state proteasomal activity of WT, and Hsp90α and Hsp90β KO cells. (K) Impacts of GA and MG132 on the proteasomal activity of an extract of WT cells measured by *in vitro* proteasomal activity assay (n=2). Proteasomal activities were measured at 20 min and 40 min after initiation of the reaction with suc-LLVY-AMC. Data are represented as total AMC fluorescence, and MG132 serves as a positive control for proteasomal inhibition. (L) Rate of proteasomal activity of extracts from WT and *Δsti1* yeast cells (BY4741 strain background) using activity reporter suc-LLVY-AMC corresponding to the steady-state proteasomal activity assay shown in Figure 3J.

**Figure S4.**
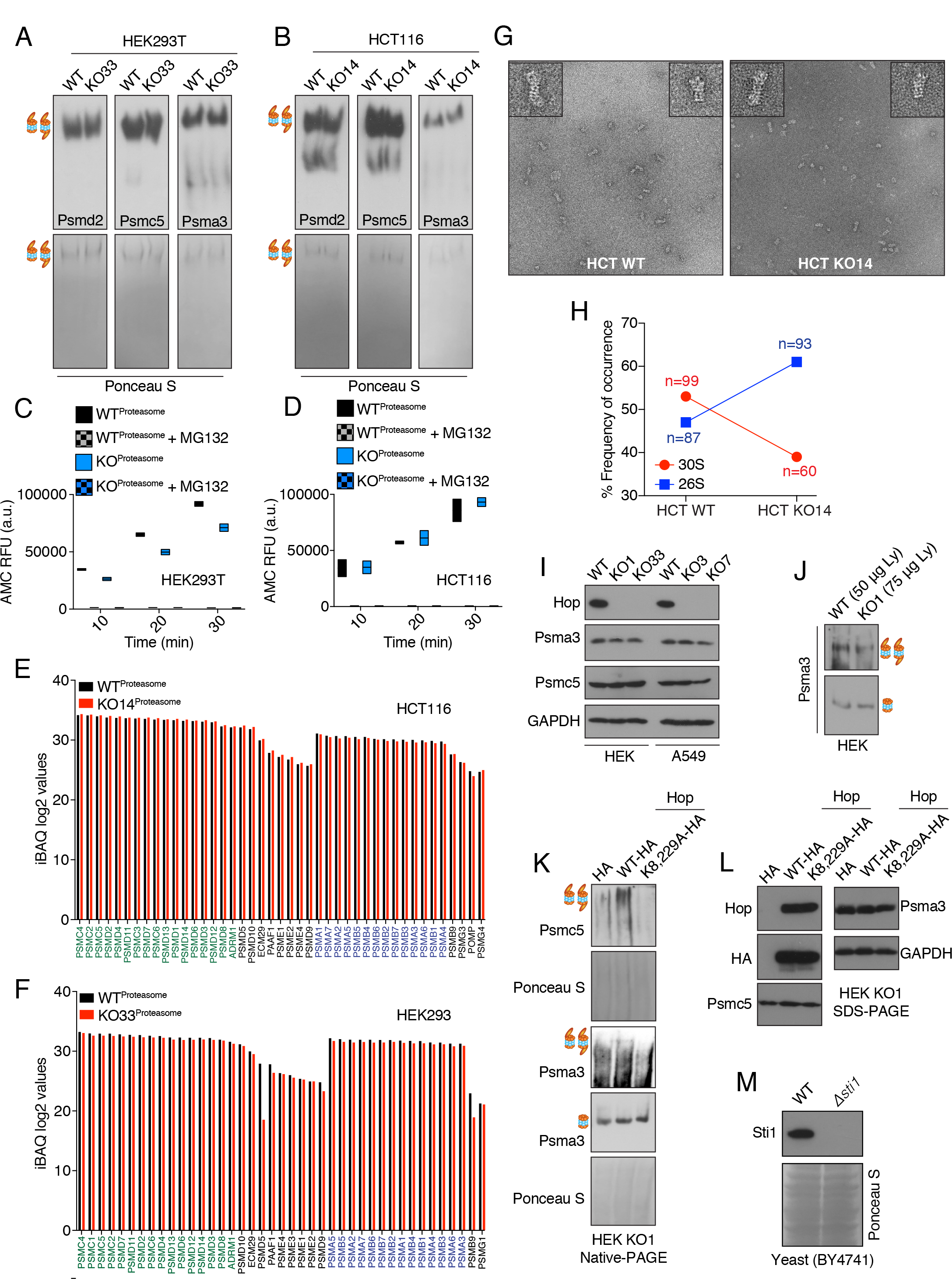
The Hsp70-Hop-Hsp90 Ternary Complex Facilitates Assembly of the Proteasome, Related to Figure 4. (A, B) Visualization of the purified 26S/30S proteasome by 4% native-PAGE. Immunoblots were probed with antibodies to Psmd2 or Psmc5 for the RP, and to Psma3 for the CP. Ponceau S stained nitrocellulose filters serve as controls of equal loading of purified proteasomes. (C, D) The activity of the purified proteasomes determined with activity reporter suc-LLVY-AMC (n=2). Data are represented as total AMC fluorescence, and MG132 serves as a positive control for proteasomal inhibition. Data are represented as a box plot. (E, F) Abundance (log2 of iBAQ values) of all proteasomal proteins identified by the MS analysis of purified proteasomes. Stoichiometric RP- and CP-specific proteasomal proteins are in green and blue, respectively. (G) Images of purified proteasome particles from WT and KO14 HCT116 cells obtained by negative staining TEM. Insets of each micrograph: side view of double-capped 30S (top left) and single-capped 26S (top right) proteasome particles. (H) Frequencies of side views of proteasomal structures. n represents the total numbers of structural projections of proteasome particles analyzed by TEM. (I) Immunoblots of the endogenous Psmc5 and Psma3. The same whole cell extracts were used for the native-PAGE analysis of Figure 4G-H. (J) Abundance of different proteasomal complexes determined by 4% native-PAGE. Note that 1.5-fold more cell lysate (Ly) of KO than of WT cells was loaded for this experiment. Antibodies to Psma3 were used to probe for proteasomal complexes. (K) Native-PAGE and subsequent immunoblot analyses of extracts from KO cells overexpressing exogenous WT and TPR double mutant (K8,229A) Hop. Antibodies to Psmc5 and Psma3 were used to probe for proteasomal complexes. An extract of KO cells transected with empty vector was used as a negative control. (L) Control immunoblot analyses of the same cell extracts of panel K. (M) Immunoblot of Sti1 protein level in WT and *Δsti1* yeast cells.

**Figure S5.**
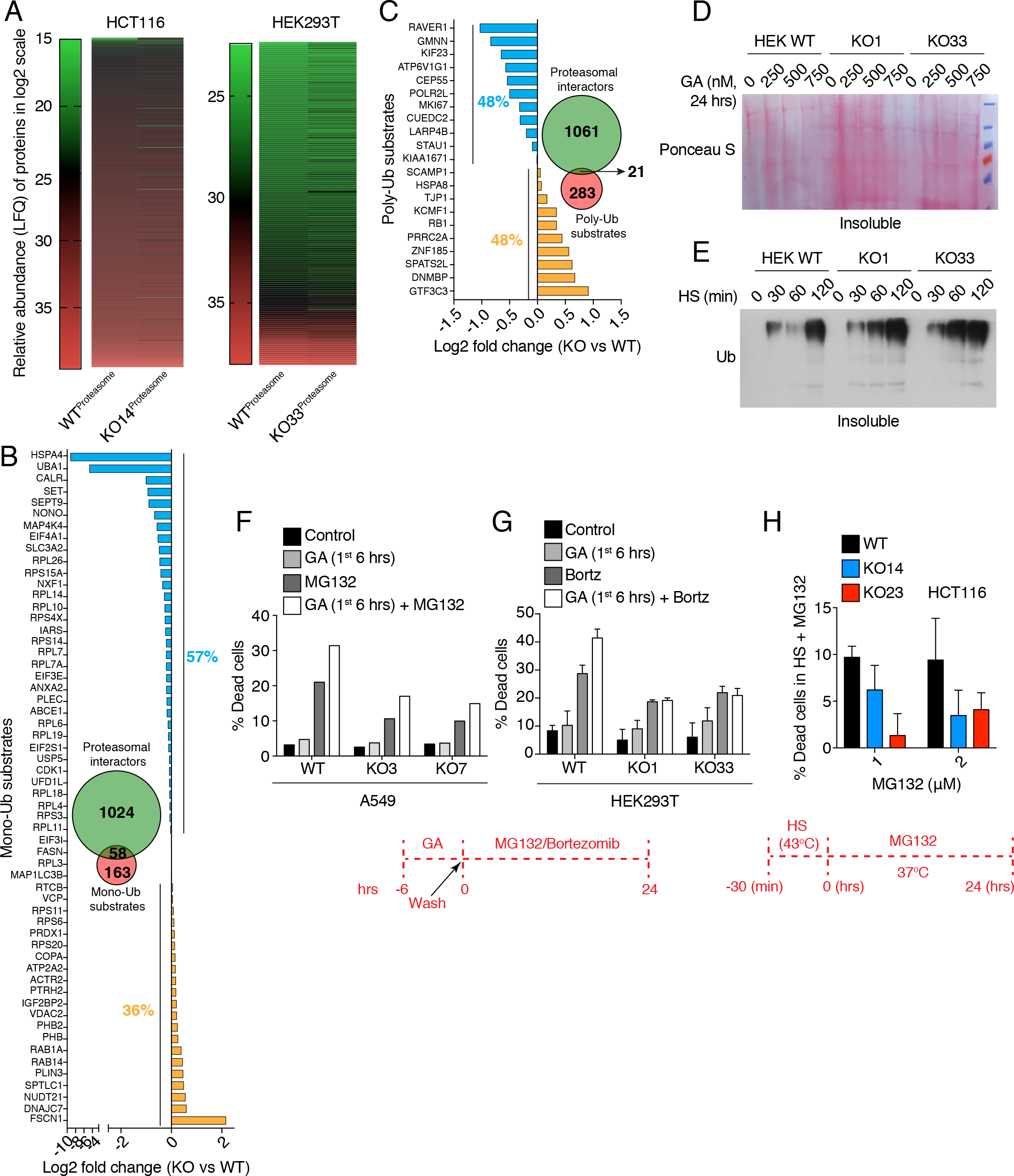
Proteasomal Degradation Flux and Utilization Are Reduced in KO Cells, Even in Proteotoxic Stress, Related to Figure 4. (A) Heat maps of the proteins associated with the purified proteasome. The proteasomal core components themselves are not included. Scale bars represent the log2 of the LFQ values of the MS analyses. (B, C) Normalized fold changes of abundance (LFQ values) of potential mono-ubiquitinated (panel B) and poly-ubiquitinated (panel C) substrates (Braten et al., 2016) identified by the MS analysis of proteasomes purified from HCT116 cells. The colors blue and yellow in the bar graphs indicate substrates, which are more and less abundant, respectively, in proteasome preparations from WT cells. Insets: Venn diagrams showing the number of proteins from our proteasome MS analysis except for the proteasomal core components themselves (in green) and proteasomal substrates from a previously published dataset (in red) (Braten et al., 2016); the respective bar graphs only show the proteins in the intersect. (D) Nitrocellulose filter stained with Ponceau S displaying the GA-induced aggregation of proteins. (E) Immunoblot of the insoluble ubiquitinated proteins after a 43°C HS. The loaded material is from equal numbers of cells. (F, G) Flow cytometric quantification of cell death induced by MG132 (10 μM, n=2; panel F) and bortezomib (500 nM; panel G) alone and in combination with a 6 h pretreatment with GA (500 nM). (H) Flow cytometric quantification of cell death induced by treatment with MG132 for 24 hrs during the recovery phase at 37°C of a HS (43°C, 30 min).

**Figure S6.**
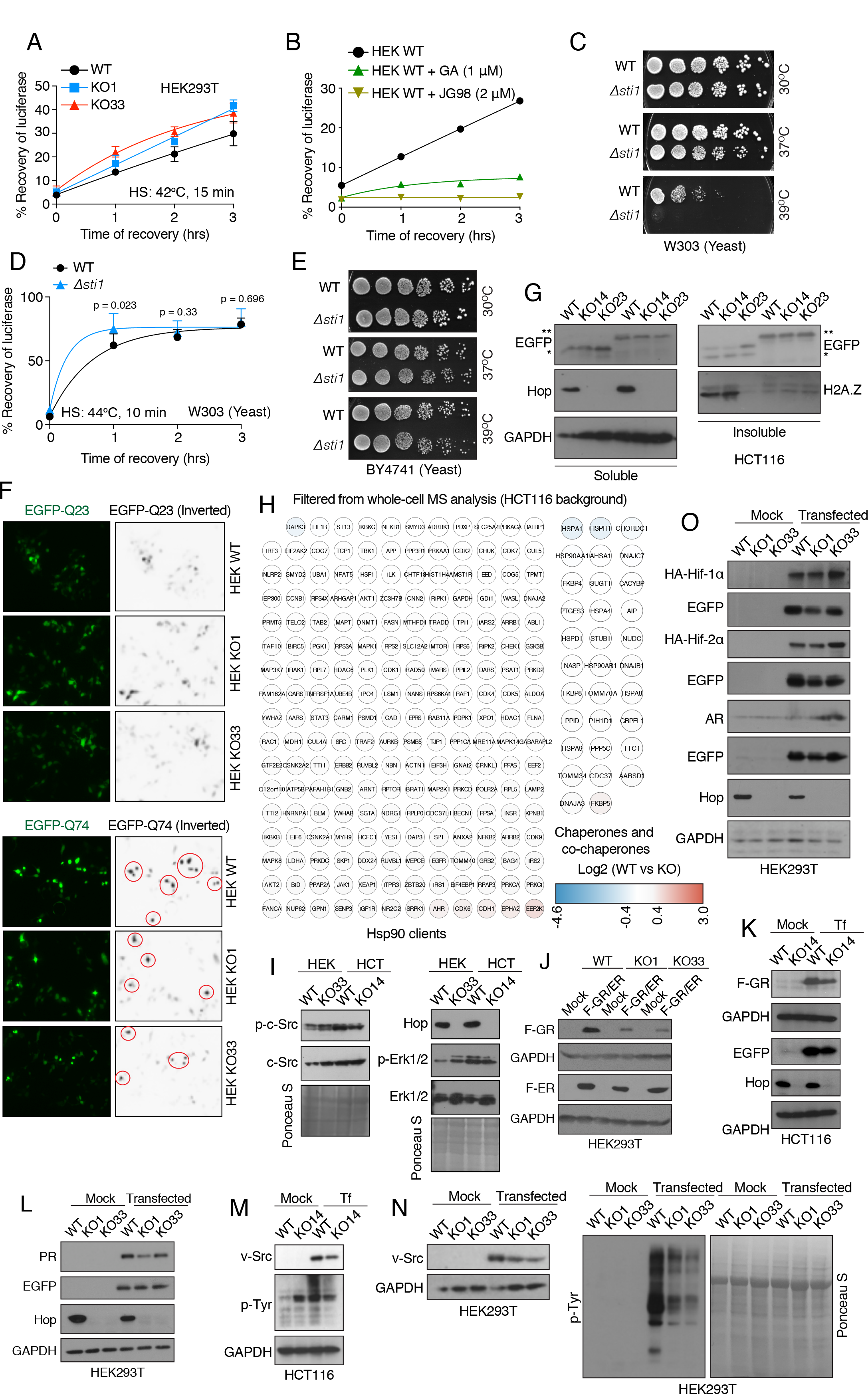
Hop-independent Enhanced Chaperoning by Hsp70 and Hsp90 in Human and Yeast Cells, Related to Figures 5. (A) *In vivo* refolding of heat-denatured luciferase. (B) *In vivo* luciferase refolding in WT HEK293T cells treated with GA (1 *μ*M) and JG-98 (2 *μ*M) (n=2) during the recovery phase at 37°C. (C) Growth assays with WT and *Δsti1* yeast cells of the W303 strain background. Spot tests of serial 5-fold dilutions; cells grown at indicated temperatures for 3 days. (D) *In vivo* refolding of heat-denatured luciferase in WT and *Δsti1* yeast cells of the W303 strain background during the recovery phase at 300C. (E) Growth assays with yeast cells of the BY4741 strain background as in panel C. (F) Fluorescence micrographs of cells expressing the fusion proteins Q74-EGFP and Q23-EGFP. Q74-EGFP aggregates are visible as punctate green fluorescence. Aggregates appear as dark black dots marked with red circles in the inverted grayscale images. The non-aggregating Q23-EGFP serves as a negative control. (G) Solubility of aggregation-prone polyglutamine model protein Q74-EGFP in WT and KO HCT116 cells. Immunoblots as in Figure 5G. (H) Heat maps of the normalized fold changes of the levels of Hsp90 clients (left), and molecular chaperones and co-chaperones (right) identified by whole cell MS analyses. The scale bar represents the log2 fold changes (WT vs KO) of the LFQ values. Immunoblots of c-Src (left part) and (I)Erk1/2 (right part). Nitrocellulose filters stained with Ponceau S indicate equal protein loading. (J) Immunoblots of F-GR and F-ER overexpressed in HEK293T cells. (K) Immunoblots of F-GR overexpressed in HCT116 cells. Tf, transfected cells. (L) Immunoblots of PR overexpressed in HEK293T cells. (M-N) Accumulation and activity of overexpressed v-Src in the indicated cellular backgrounds. Tyrosine-phosphorylated total protein is indicative of v-Src activity. The nitrocellulose filter stained with Ponceau S serves as a loading control. Tf, transfected cells. (O) Immunoblots of overexpressed HA-tagged Hif-1α and Hif-2α, and of androgen receptor (AR). EGFP serves as a transfection control.

**Figure S7.**
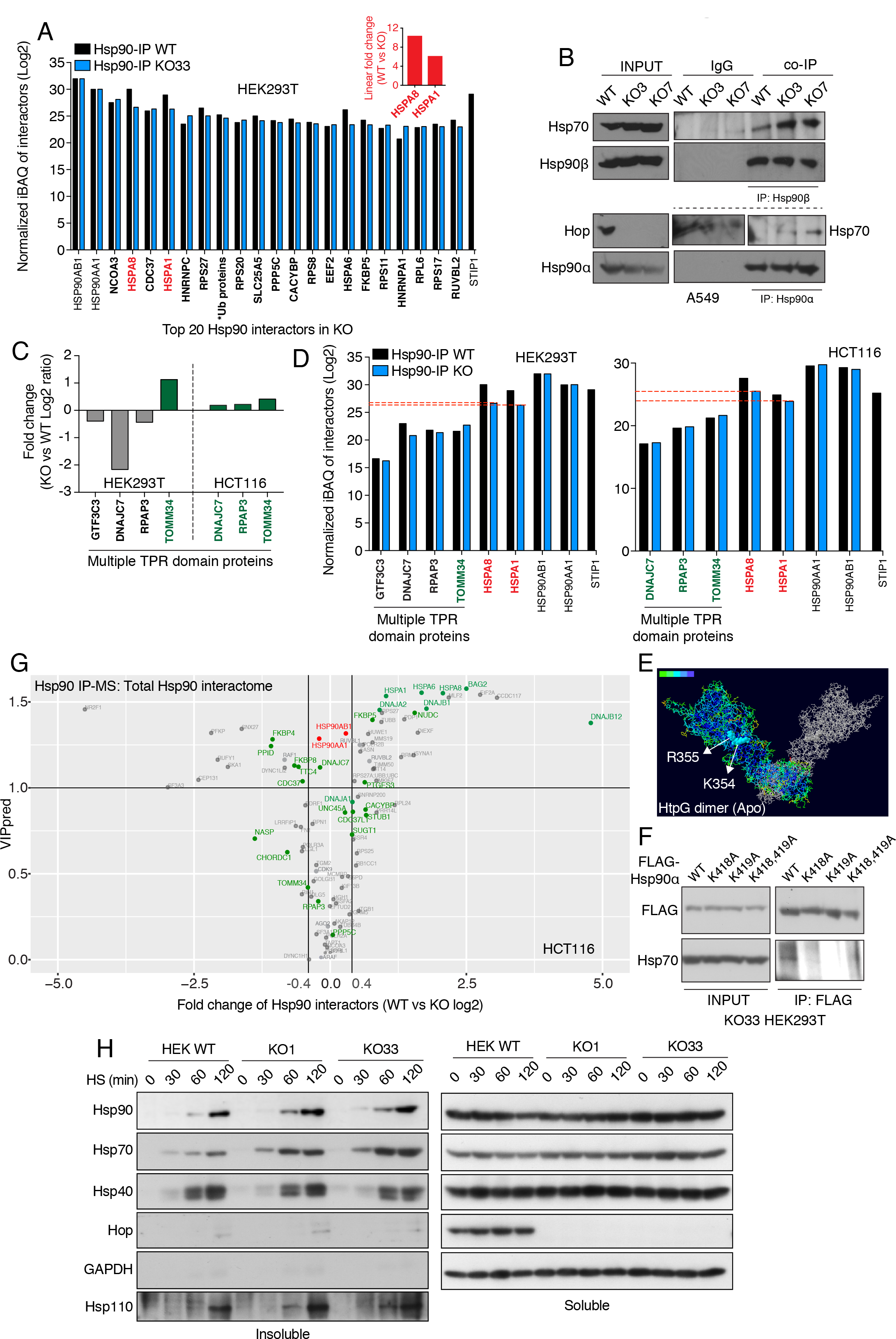
Human Hsp70 and Hsp90 Interact Directly and Form a Functional Prokaryote-like Molecular Chaperone Complex Even in the Absence of Hop, Related to Figures 6 and 7. (A) Abundance of the top 20 Hsp90 interactors (according to the highest iBAQ values) of KO HEK293T cells (names in bold). The graph shows the iBAQ values as log2 of the Hsp90-IP-MS analyses. Hsp90α (HSP90AA1) and Hsp90β (HSP90AB1) were the bait proteins, and absence of Hop (STIP1) serves as quality control marker for KO cells. Inset: Linear fold changes of the values for Hsp70 (HSPA1) and Hsc70 (HSPA8). *Ub proteins: UBB, UBC, UBA52, RPS27A. (B) *In vivo* interaction of Hsp90 and Hsp70 in A549 cells as determined by an immunoprecipitation experiment. (C) Bar graph showing the enrichment (green bars) or depletion (grey bars) of multiple TPR domain-containing proteins identified in the Hsp90-IP-MS analyses. (D) Relative abundance of the multiple TPR domain-containing proteins compared to Hsc70 (HSP8) and Hsp70 (HSPA1). The abundance of both Hsp70 and Hsc70 is indicated by red dotted lines. The graph shows the iBAQ values as log2 of the Hsp90-IP-MS analyses. (E) Surface accessibility of the highlighted amino acids in the dimeric structure of HtpG in its open conformation. Color code as in Figure 6F. (F) Interaction of Hsp90 point mutants with endogenous Hsp70. Immunoprecipitation of exogenously overexpressed FLAG-tagged WT and point mutant Hsp90α from KO cells. Immunoblots were probed with antibodies to FLAG or Hsp70. (G) The volcano plot represents the normalized fold changes of the Hsp90 interactors identified by the Hsp90-IP-MS analysis with HCT116 cells versus their VIP values derived from the OPLS-DA model. Cut-offs and color as in Figure 7A. (H) Immunoblots showing time-dependent HS-induced recruitment of Hsp90, Hsp70, Hsp40 and Hsp110 into insoluble protein aggregates. The soluble protein fractions show the total levels of the aforementioned proteins. GAPDH serves as a control for soluble proteins even following HS. The loaded material is from equal numbers of cells. HS was at 43°C for the indicated time points. Note that the weak bands of the 120 min time points in the Hop immunoblots are non-specific.

